# Local and long-distance inputs dynamically regulate striatal acetylcholine during decision making

**DOI:** 10.1101/2022.09.09.507130

**Authors:** Lynne Chantranupong, Celia C. Beron, Joshua A. Zimmer, Michelle J. Wen, Wengang Wang, Bernardo L. Sabatini

## Abstract

Within the basal ganglia, striatal dopamine (DA) and acetylcholine (Ach) are essential for the selection and reinforcement of motor actions and decision making. *In vitro* studies have revealed a circuit local to the striatum by which each of these two neurotransmitters directly regulates release of the other. Ach, released by a unique population of cholinergic interneurons (CINs), drives DA release via direct axonal depolarization. In turn, DA inhibits CIN activity via dopamine D2 receptors (D2R). Whether and how this circuit contributes to striatal function *in vivo* remains unknown. To define the *in vivo* role of this circuit, we monitored Ach and DA signals in the ventrolateral striatum of mice performing a reward-based decision-making task. We establish that DA and Ach exhibit multiphasic and anticorrelated transients that are modulated by decision history and reward outcome. However, CIN perturbations reveal that DA dynamics and reward-prediction error encoding do not require Ach release by CINs. On the other hand, CIN-specific deletion of D2Rs shows that DA inhibits Ach levels in a D2R-dependent manner, and loss of this regulation impairs decision-making. To determine how other inputs to striatum shape Ach signals, we assessed the contribution of projections from cortex and thalamus and found that glutamate release from both sources is required for Ach release. Altogether, we uncover a dynamic relationship between DA and Ach during decision making and reveal modes of CIN regulation by local DA signals and long-range cortical and thalamic inputs. These findings deepen our understanding of the neurochemical basis of decision making and behavior.

## Introduction

The basal ganglia are a group of interconnected sub-cortical nuclei that integrate information from cortex, thalamus, and mid-brain dopamine centers, amongst others, to modulate goal directed behavior, including motor learning, habit formation, and decision making^1–3^. The striatum is the principal input structure of the basal ganglia, and its function is controlled by a complex array of neurotransmitters and neuromodulators arising from both local and distant sources^4–7^. Among these is dopamine (DA), which is released in striatum by long-range axons arising from midbrain ventral tegmental area (VTA) and substantia nigra pars compacta (SNc) neurons^8–11^. DA neurons (DANs) drive reinforcement learning by encoding reward prediction error (RPE), the difference between experienced and expected reward, and, mechanistically, by regulating aspects of neuronal and synapse function^12–16^. Disruption of DA signaling contributes to many debilitating psychomotor disorders, including Parkinson’s disease and drug addiction, underscoring the significance of this neurotransmitter in striatal function^17–19^.

In addition to having the highest concentrations of DA and DA receptors in the mammalian brain, the striatum also contains some of the highest levels of acetylcholine (Ach)^20,21^. Ach is primarily released by local cholinergic interneurons (CINs), a specialized and rare cell type that comprises only 1-2 % of striatal neurons but that is tonically active and makes dense and extensive intra-striatal axonal arborizations^22,23^. Pioneering studies in primates revealed that CINs characteristically reduce or ‘pause’ their firing in response to both appetitive and aversive stimuli ^23–28^. These responses emerge over the course of learning, leading to the hypothesis that CINs, like DANs, modulate reinforcement learning and goal-directed behaviors. CIN pauses in turn may alter the plasticity of corticostriatal synapses to support procedural learning ^29^, but a full understanding of how CINs contribute to striatal function is lacking.

Bidirectional interactions between DA and Ach release have been observed within striatum. In Parkinson’s disease, loss of DA leads to a hypercholinergic state, and anti-cholinergic drugs were amongst the earliest treatments that alleviated symptoms of this disease^30,31^. During learning, increases in DAN activity coincide with CIN pauses, which are abolished upon DAN lesions^23,24^. Studies *in vitro* uncovered a local circuit by which DA and Ach directly influence each other. Synchronized firing of multiple CINs activates nicotinic acetylcholine receptors that are located on and depolarize DAN axons^32–35^. If of sufficient amplitude, this axonal depolarization can induce a propagating axonal action potential that evokes DA release within striatum (**Fig. 1a**)^35^. In turn, DA potently inhibits CIN activity by acting on Type-2 DA receptors (D2Rs) expressed by CINs (**Fig. 1a**)^36–38^.

**Fig. 1.**
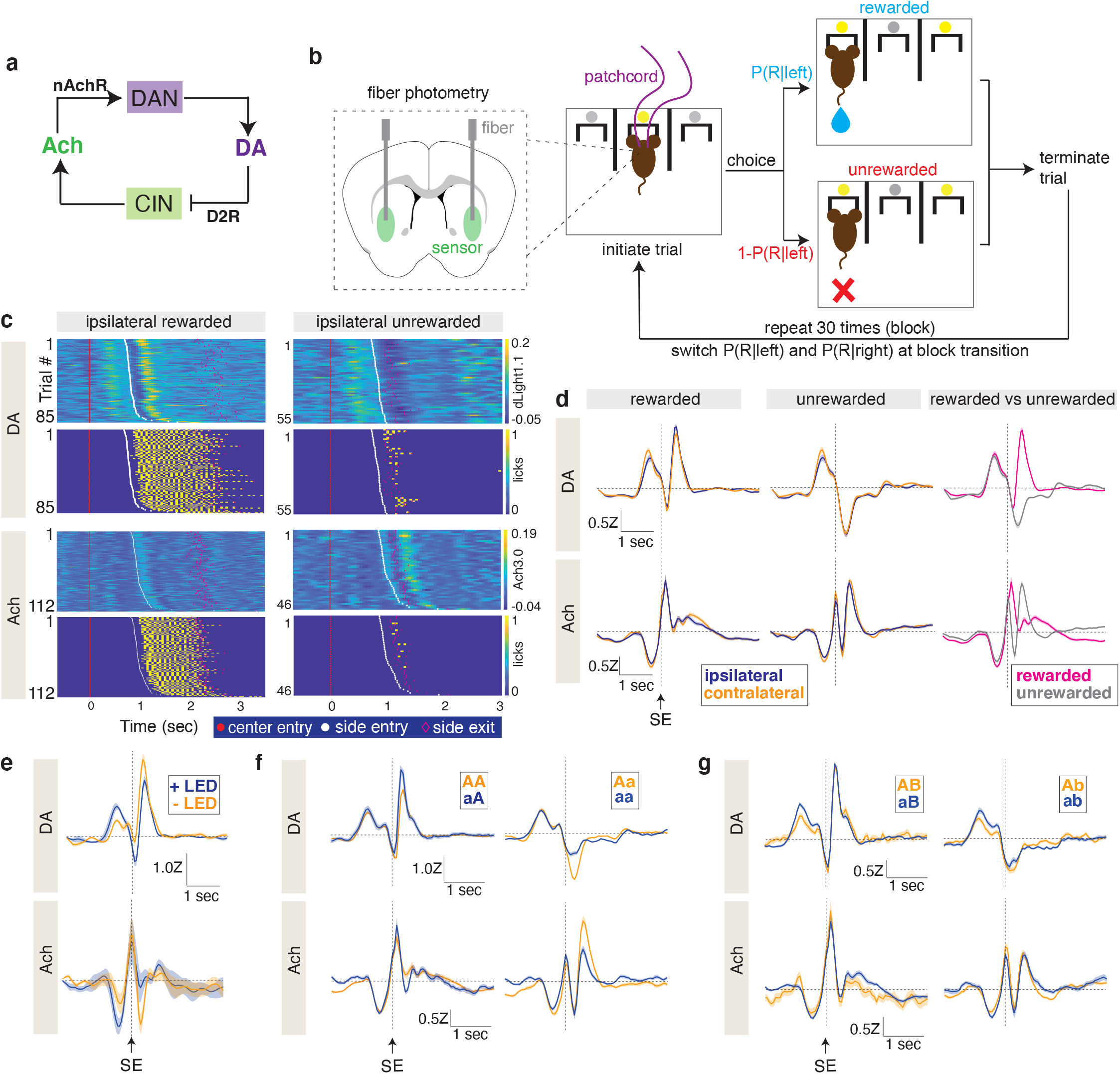
Multiphasic dynamics of dopamine and acetylcholine in the VLS during reward-based decision making. **a**. Schematic of proposed interactions between dopamine and acetylcholine within the striatum. Cholinergic cells (CINs) release acetylcholine (Ach), which can evoke dopamine (DA) release via nicotinic acetylcholine receptors (nAchR) expressed on dopamine neuron (DAN) terminals. Conversely, DA inhibits CINs via dopamine type 2 receptors (D2Rs). **b**. Schematic of the 2ABT. An LED (yellow circle) above the center port cues the mouse that a trial can be initiated. Following a center port poke to initiate a trial, the mouse makes a choice to go to the left or right port. The reward probability is different for each port and used to probabilistically deliver reward to the side-port chosen by the mouse. The trial is terminated when the mouse exits the side port such that only one port can deliver reward per trial. Fiber photometry recordings are performed continuously during the task from the ventrolateral striatum of mice expressing a fluorescent sensor (left dashed inset). **c**. Ipsilateral DA (dLight1.1) and Ach (Ach3.0) dynamics (color table) and associated licks (yellow) recorded from an example mouse during a 2ABT session. Each row shows data from an individual trial aligned to center entry (red dots), sorted by increasing side port entry time (white dots), and with side port exit time denoted (pink diamonds). **d**. DA and Ach release in ipsilateral and contralateral trials (left and middle), as well as for rewarded and unrewarded trials in which ipsilateral and contralateral trials are combined (right). The averaged Z scored sensor signal for the indicated trial type (bold line) is depicted with the standard error of the mean (SEM) (shaded region). Data are aligned to side port entry (SE) (DA: n = 7 mice; Ach: n = 9 mice). **e**. DA and Ach release during rewarded and unrewarded trials with omission of the LED cue during the 2ABT. Data is depicted as in (**d**) (n = 4 mice). **f**. DA and Ach release for trials in which the mice choose the same side in both the current and previous trials, segregated by the indicated choice and reward outcome histories. ‘A’ indicates a rewarded trial and ‘a’ indicates an unrewarded trial. Data are depicted as in (**d**) (DA: n = 7 mice; Ach: n = 9 mice). **g**. DA and Ach release for trials in which the mice choose the opposite side as the previous trial, segregated by the indicated choice and reward outcome histories. ‘B’ and ‘b’ are rewarded and unrewarded trials, respectively, performed on the side opposite to that of ‘A’ and ‘a’. Data are depicted as in (**d**) (DA: n = 7 mice; Ach: n = 9 mice).

Despite a detailed understanding of the mechanisms of interaction between DA and Ach *in vitro*, if, when, and how these control DA and Ach levels to regulate striatal function *in vivo* are unknown. In terms of CIN-mediated control of DAN axons, it is unclear if sufficient CIN synchronization occurs and to what degree nicotinic Ach receptors are available to evoke DA release *in vivo*^39^. Furthermore, it is not known how much and when the potential CIN influence on DA signaling is functionally significant compared to the direct somato-dendritic modulation of DAN activity. The ability for CINs to evoke DA release independently of somatic firing has been proposed as a mechanism to explain differences between DAN somatic activity and striatal DA levels^40^; however, recent studies report robust correlation between somatic and axonal signaling^41,42^. In terms of DA-mediated inhibition of CINs, although CIN pauses can be induced by activation of D2Rs, they can also be triggered by other striatal inputs, including cortical and thalamic glutamatergic projections and long-range GABAergic VTA inputs^43–45^. Indeed, CIN-specific deletion of the gene encoding D2Rs, *Drd2*, reduces, but does not abolish, the CIN pause in a reward-based task, suggesting additional sources are responsible^46^.

To provide a more complete understanding of what factors shape Ach release during reward-based decision making and how Ach release contributes to DA release, we examined striatal Ach and DA levels during a complex reward-based decision-making task in mice. We find that behavioral events drive multiphasic and anticorrelated responses in DA and Ach within striatum, and the history of choice and reward outcome alters the release of both neurotransmitters. Surprisingly, elimination of Ach release from CINs has minimal effects on DA dynamics during the task, suggesting that DA release in this context is largely insensitive to local, CIN-dependent regulation. In contrast, striatal Ach levels are shaped directly by local DA levels: DA reduces Ach levels in a D2R-dependent manner, and disrupting this regulation impairs performance of mice in the task. Finally, neurotransmitter release from cortical and thalamic axons are necessary for normal striatal Ach dynamics and levels. These results show that local striatal interactions between Ach and DA are neither fixed nor always dominant in driving the dynamics between the neuromodulators during decision making. Instead, DA modulates Ach levels only at specific times. Thus, the transients and resulting relationship between these two neuromodulators arise from a combination of intra- and extra-striatal inputs.

## Results

### Striatal Ach and DA levels are dynamically regulated during reward-based decision making

Decision making in probabilistic reward-reinforced tasks requires that individuals integrate internal and external cues in the context of a changing environment to flexibly make choices that maximize desired outcomes. To examine the local circuit interactions between striatal Ach and DA (**Fig. 1a**) during, and their contributions to, such behaviors, we monitored neuromodulator levels in mice performing a dynamic and probabilistic two-port choice task modeled after paradigms that engage striatal pathways and require striatal activity for optimal performance^47–49^. In this two-armed bandit task (2ABT), mice move freely within a box containing three ports (**Fig. 1b**). An LED cue above the center port signals that a mouse can initiate a trial by placing its snout (i.e., “poking”) in the center port. The mouse must then choose to poke into either a left or right port, each of which probabilistically delivers water after snout entry. In a block structure (30 rewards between block transitions) either the left or right port is designated as “high reward probability” (*P(reward)=P*_*high*_) and the other port as “low reward probability” (*P(reward)=1-P*_*high*_). To efficiently obtain rewards, the mouse must learn which is the high reward probability port in that block and detect when block transitions occur. This task structure requires mice to employ flexible decision-making strategies and integrate information about previous trial outcomes to sample and choose ports.

Mice become proficient in this task and robustly alter their port selections at block boundaries (**Extended Data Fig. 1a**). In the middle of blocks, expert mice typically repeatedly choose the highly rewarded port and occasionally sample the low-reward probability port (**Extended Data Fig. 1a**). However, following successive unrewarded trials resulting from reward probability reversals at block transitions, they transiently increase their probability of switching ports between trials and rapidly change their choice to the new high reward probability port (**Extended Data Fig. 1c and 1d**). As a result of these strategies, proficient mice achieve highly similar decision times, reward rates, and switching rates (**Extended Data Fig. 1b**). During this behavior we capture the timing of port entries and withdrawals, the timing and number of licks at each port, and the trial outcomes. Center port entry and exit occur in rapid succession, followed by a delayed entry into the side port (**Extended Data Fig. 1e**). In rewarded trials, the water reward is triggered by side port entry, and mice repeatedly lick to consume the reward whereas in unrewarded trials the mice rarely lick the port (**Fig. 1c, Extended Data Fig. 1e third column**). Mouse behavior and evidence accumulation in the task can be summarized by a variety of models^49–51^. Here, we will employ a recursively-formulated logistic regression (RFLR, see methods) that was developed from a behavior task similar to the one we employ^49^. The RFLR model utilizes three parameters to capture, respectively, the tendency of an animal to repeat its last action (alpha, *α*), the relative weight given to information about current action and reward (beta, *β*), and the time constant over which action and reward history decay (tau, *τ*). The RFLR model accurately captures the switching probability across multiple choice and outcome histories (**Extended Data Fig. 1g**) as well as the switching dynamics at block transitions (**Extended Data Fig. 1h**). RFLR coefficients are also comparable across expert mice (**Extended Data Fig. 1i**). Overall, mice achieve high proficiency on a probabilistic reward task, and their behavior can be accurately captured by a reduced logistic regression model.

To determine how DA and Ach signals change during this behavior, we used frequency-modulated fiber photometry to record the fluorescence of the genetically-encoded sensors for DA (dLight1.1 ^52^ and Ach (GRAB-Ach3.0, abbreviated as Ach3.0)^53^ expressed in separate hemispheres within the ventrolateral portion of the dorsal striatum (VLS) (**Extended Data Fig. 2a**), a region previously associated with controlling the behavior of mice in reward-based decision-making tasks^54^. Fluorescence transients were frequency-demodulated, down-sampled to 54 ms time bins (see Methods), and Z-scored over a 60 second rolling window. We observed robust and multiphasic DA and Ach transients in individual trials that differed depending on rewarded outcome (**Fig. 1c and 1d**). These transients reflect a combination of the release, reuptake, and degradation of the neuromodulator. Importantly, they are not observed in ligand-binding site mutants of the sensors expressed in VLS and recorded within the same behavioral context (**Extended Data Fig. 2b and 2c**), confirming their dependence on neuromodulator binding. Despite the lateralization of the task, the DA and Ach transients are similar for trials in which the mouse selected the port ipsilateral versus contralateral to the fiber placement (**Fig. 1d, left and middle**); therefore, we combine data across both trial types (**Fig. 1d, right)**.

To understand what behavioral features impact DA and Ach transients, we compared their profiles during rewarded and unrewarded trials. As expected, DA signals changed at the instances of task-relevant behavioral events, such as during movement from the center port to the side port (**Fig. 1c and 1d**). Furthermore, DA levels diverge depending on reward outcome, rising in rewarded trials but falling during unrewarded trials (**Fig. 1c and 1d**), consistent with encoding of RPE.

In contrast to past reports that striatal Ach is outcome-insensitive^25^, we found that reward outcome clearly modulates Ach transients. However, unlike DA, the sign of Ach transients is not strictly opposite in rewarded versus unrewarded trials (**Fig. 1d, lower right**). Further support for reward-outcome modulation of both DA and Ach is revealed during trials in which the LED cue that signals trial initiation is omitted. In the absence of this cue, the mice can still obtain rewards, but they are more unexpected. Compared to control rewarded trials, the DA transient in cue-omission rewarded trials is smaller during the center to side port transition but is larger when reward is delivered (**Fig. 1e**). Like DA, Ach transients are also affected by cue omission, although generally in the opposite direction to DA transients: the magnitude of the Ach transient is increased during center-to-side transition and decreased during reward acquisition.

Segregating the trials by reward outcome alone masks the complex and interrelated effects that choice and reward history can have on behavior and neurotransmitter release. Therefore, we subdivided rewarded and unrewarded trials by the task history and find that the outcome on the previous trial strongly modulates outcome-dependent DA and Ach levels on the current trial. To concisely represent choice and reward history, we use previously established annotation in which each trial is assigned a letter label that captures both the chosen action and the outcome: the letter identity signifies the port choice and the capitalization signifies outcome^49^. Specifically, ‘A’ and ‘a’ represent rewarded and unrewarded trials on one side, respectively, whereas ‘B’ and ‘b’ represent rewarded and unrewarded trials, respectively, on the alternate side. In a multi-trial sequence, the port selected in the first trial is designated a/A. For example, in a two-trial sequence, aB denotes an unrewarded trial in the first port (‘a’) followed by a rewarded trial in the other port (‘B’).

When a mouse chooses the same port in two consecutive trials, DA signals increase more and Ach signals decrease more following side port entry for a rewarded trial that was preceded by an unrewarded trial (aA) rather than a rewarded trial (AA), reflecting different reward expectations due to previous experience (**Fig. 1f, left**). Similarly, for an unrewarded trial, the magnitudes of the dip in DA levels and the concomitant rise in Ach levels are greater if it was preceded by a rewarded trial (Aa) rather than by an unrewarded trial (aa) (**Fig. 1f, right**). Interestingly, these effects of expectation on trial-outcome DA and Ach are absent if the mouse switches ports between trials: i.e., the signals following side-port entry are similar for AB and aB trials and similar for ab and Ab trials (**Fig. 1g**). However, on these selection-port switch trials, the DA signals associated with the center-to-side port transition were greater when the previous trial was unrewarded (**Fig. 1g**). Thus, when a mouse makes the decision to switch ports from one trial to the next, it approaches this choice in a different state shaped by history; however, any outcome resulting from this switch is equally unexpected.

Across multiple instances in which reward expectation modulates both DA and Ach transients, we observed that changes in DA and Ach are often temporally coincident but in the opposite direction. This is in line with previous observations in primates that CIN pauses are time-locked with increases to DAN activity^25^. This apparent negative correlation may arise from a direct interaction between DANs and CINs (i.e. DA mediated inhibition of CIN activity; **Fig. 1a**) or, alternatively, point to opposite modulation of both signals by a common input. Nevertheless, during the task, the relationship between DA and Ach is neither simple nor fixed as there are periods in which both signals go up or down synchronously or independently, suggesting a flexible and dynamic coupling between the two neuromodulators.

### DA and Ach dynamics are driven by multiple behavioral events and action-outcome history

To formally and quantitatively evaluate the contribution of behavior events to DA and Ach levels, we developed a general linear model (GLM) to predict neuromodulator levels from behavior (**Extended Data Fig. 3a**). In the simplest model, which we term the ‘base GLM’, we incorporated predictive variables based on key behavioral events: center port entry and exit, side port entry and exit, reward delivery, licks, and side port entries that occur outside of the trial structure. For each event, the model derived kernels are comprised of time shifted *β* coefficients that minimize an ordinary least squares (OLS) cost function. These kernels are convolved with impulse functions representing the timing of behavior events and summed to reconstruct the photometry signal (**Extended Data Fig. 3a**). Regularization was not required for this base GLM as training, validation, and test datasets yielded similar MSE, and incorporation of L1 and/or L2 regularization did not improve model performance compared to OLS (**Extended Data Fig. 3b**); thus, the relatively few model parameters do not lead to overfitting. We find that this simple model captures most variance across the trial-associated data (DA GLM R^2^ = 0.206; Ach GLM R^2^ = 0.206) (**Extended Data Fig. 3c and 3d, right**).

To assess the contribution of each behavioral variable to GLM performance, we performed a ‘leave-out analysis’ of individual features (**Extended Data Fig. 3e**). For both DA and Ach models, center entry and center exit are likely redundant as loss of either does not impact model fit. Meanwhile, omission of only side entry and reward variables greatly weakens the GLM performance for DA signals. In contrast, loss of side entry, reward, side exit and lick inputs all impair the GLM performance for Ach signals. These differences highlight the unique influence of each behavioral event on DA and Ach dynamics.

Although the base GLM robustly captures the true signals of both Ach and DA, there are discrepancies with the reconstructed signals across several histories (**Extended Data Fig. 3c and 3d, right**). Given the importance of choice and reward history for modulating the signals of both neurotransmitters (**Fig. 1f and 1g**), we expanded the feature set of the base GLM to include side port entries segregated by the eight possible action-outcome combinations, which we term the ‘history GLM’ (**Extended Data Fig. 4a and 4b)**. Indeed, inclusion of these parameters reduced the MSE between the predicted and test data for both DA and Ach GLMs (**Extended Data Fig. 3e, + history MSEs**). This reflects an improvement in the ability of the history GLMs to capture the variance of the trial-associated data without overfitting (**Extended Data Fig. 3e**) (DA GLM R^2^ = 0.213; Ach GLM R^2^ = 0.214). Furthermore, despite the addition of multiple parameters, regularization was still not required for optimal model performance (**Extended Data Fig. 4c**). Altogether, by modeling DA and Ach signals with GLMs, we reveal the influence of multiple behavioral variables and action-outcome history on the dynamics of both neurotransmitters during decision making.

### Ach and DA levels are anticorrelated across trial types, except at moment of choice-evaluation

Because DA and Ach may directly interact *in vivo*, we characterized the relationship between their patterns of release to determine if they support either the positive or negative interactions proposed. Comparison of the trial-averaged fluorescence signals and the GLM analyses suggest that such interactions exist: an overlay of DA and Ach dependent signals recorded from separate hemispheres reveals a striking reciprocal relationship between the two neurotransmitters during rewarded and unrewarded trials (**Extended Data Fig. 5a**). Nevertheless, following side port entry of unrewarded trials, DA dips while Ach rises twice (**Extended Data Fig. 5a, orange arrows)** which suggests a complex and potentially dynamic relationship between the two neurotransmitters (**Extended Data Fig. 5a, right**). Cross correlation analysis of these trial-segregated signals in which DA lags Ach reveals an anticorrelation that occurs with a positive time lag (**Extended Data Fig. 5b**), indicating that changes in DA are associated with delayed changes in Ach of the opposite sign. In addition, there is a positive correlation that occurs with a negative time lag, suggesting that changes in Ach are followed by changes in DA of the same sign.

To more accurately assess the dynamics of and relationship between DA and Ach transients, we performed simultaneous recordings of both neurotransmitters within the same hemisphere by coexpressing a red-shifted DA sensor, rDAh^55^ and the green Ach sensor (**Fig. 2b, left**). To understand the impact of switching DA sensors, we exploited the fact that the release of DA is highly correlated across hemispheres within the same mouse (**Extended Data Fig. 5c**), allowing us to directly compare DA signals detected by rDAh versus dlight1.1. Both sensors yield similar signals, albeit with a smaller amplitude for rDAh (**Fig. 2a**), likely reflecting its slower kinetics and higher affinity for DA compared to dLight1.1. Simultaneous DA and Ach recordings within the same hemisphere (**Fig. 2b**) recapitulate what we observed with recordings from separate hemispheres. The continuous, non-trial segregated DA and Ach signals are highly anticorrelated with a positive time lag (**Fig. 2c**), as are the trial-segregated DA and Ach signals (**Fig. 2d, left**) and the residuals of these transients (**Fig. 2d, right**) (i.e. signal and noise correlations, respectively). These relationships are consistent with the proposed intra-striatal interactions between Ach and DA, but they may also be independent of these direct interactions and arise from a common input that directly regulates DANs and CINs.

**Fig. 2.**
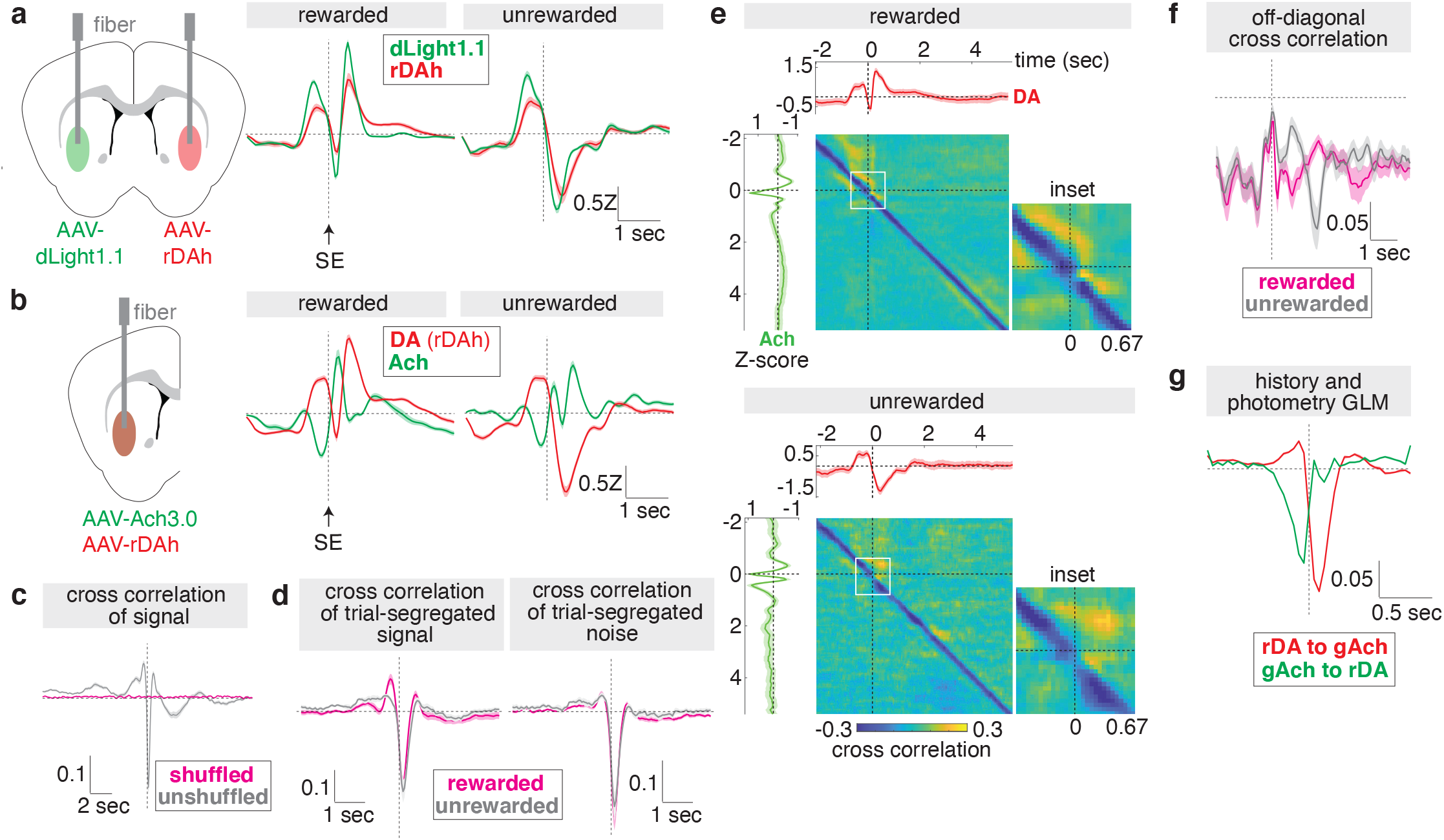
Ach and DA signals are dynamically correlated during reward-based decision making. **a**. Comparison of DA release detected by dlight1.1 and rDAh (middle and right) recorded from mice with the indicated bilateral injections (left). The average Z-scored sensor signals are shown (bold line), with standard error of the mean (SEM) overlaid (shaded region; errors are often smaller than line thickness). Data are aligned to side port entry (SE) (n = 3 mice). **b**. Overlay of DA and Ach dynamics recorded simultaneously for the indicated trial types (middle and right). Schematic of the injection strategy (left). Data are depicted as in (**a**) (n = 6 mice). **c**. Covariance of DA and Ach signals recorded across an entire behavioral session, compared to shuffled data. The average covariance is depicted (bold line) with the standard error (shaded region) (n = 6 mice). **d**. Covariance of trial-segregated DA and Ach signals (left) and their noise (right) in which DA lags Ach by the indicated time. Data are depicted as in (**c**). **e**. Full time-dependent covariance analysis of DA and Ach signals in rewarded and unrewarded trials. The average DA and Ach signals (bold lines) with SEM (shaded region) are shown within the top and left subplots, respectively. An enlarged inset of the region outlined in the white box is shown to the right of each matrix, with the time (sec) indicated on the bottom of the inset (n = 6 mice). **f**. Summary of the off-diagonal negative covariance calculated from the matrices in (**e**). **g**. Photometry kernels produced by a GLM which incorporates behavioral, history, and photometry variables. The kernels (bold lines) that predict Ach signals from rDAh signals (rDA to gAch) and DA signals from Ach3.0 signals (gAch to rDA) are shown with respect to the time shift, with the confidence interval (shaded region) overlaid (n = 6 mice).

Cross-covariance analysis, as presented above, assumes a “wide-sense stationary” process in which the mean and variance of the signal do not change over time. This permits the covariance to be expressed as a function of one variable representing a time shift between the signals. However, during behavior both the mean and variance of DA and Ach levels change dynamically. Therefore, we performed full time-dependent covariance analysis, in which we calculate how variance about the mean of DA at one time point (*t*_1_) influences Ach at another time point (*t*_2_) (see Methods) (**Extended Data Fig. 5d)**. This results in a 2-dimension function – *K*(*t*_1_, *t*_2_) (**Fig. 2e**) – characterized by a strong time-lagged negative covariance, which we call the off-diagonal (**Fig. 2f**) showing that, at most time points, changes in DA are followed by opposite sign changes in Ach approximately 100 ms later. However, the strength of the covariance varies as a function of time during the trial and, intriguingly, it nearly disappears when the animal enters the side port in both rewarded and unrewarded trials (**Fig. 2e, insets**). In addition, off band positive correlations emerge selectively as the animal approaches the side port (**Fig. 2e**). Thus, the relationship between DA and Ach is dynamic and context dependent. Taken together, these results are consistent with the proposed reciprocal interactions between DA and Ach. During most periods, DA release activates D2Rs on CINs, reducing CIN activity, and leading to a time-lagged negative correlation between these signals. At specific moments, such as when the animal approaches the side port, positive correlations consistent with Ach driven DA release appear whereas at other times the negative correlation is weakened. These dynamics might be driven by additional inputs upstream of CINs and DANs whose activity could momentarily override the direct interactions between DA and Ach.

We can further quantify the interactions between simultaneously recorded DA and Ach signals using a GLM that incorporates photometry as a predictive variable alongside the behavioral variables, such that DA dynamics can predict changes in Ach and vice versa. This model generated kernels of DA and Ach (**Fig. 2g**) which highly resemble the covariance function – both are dominated by a negative interaction occurring with opposite time shifts of approximately 100 ms. These kernels are then convolved with their respective photometry traces and added to the convolved behavioral variables to generate the predictions of Ach and DA, respectively. The photometry variable alone is insufficient to predict Ach and DA signals (**Extended Data Fig. 5e**) (DA GLM R^2^ = 0.116; Ach GLM R^2^ = 0.171), underscoring the importance of behavioral events in driving Ach and DA transients. Meanwhile, incorporation of these photometry kernels into history GLMs for DA and Ach improves model performance without overfitting (DA GLM R^2^ = 0.336; Ach GLM R^2^ = 0.388). Taken together, we establish that DA and Ach signals are anticorrelated throughout decision making, but this relationship is flexible and can be transiently disrupted, notably at the moment of choice evaluation.

### Ach release from CINs is not necessary for DA dynamics during decision-making

To determine if CINs regulate striatal DA signals during decision making, we first tested if Ach release is sufficient to modulate *in vivo* DA levels in a manner as robust as what is observed *in vitro*^32,34^. Expression of the activating opsin Chrimson^56^ in CINs in the VLS enables laser-dependent stimulation of Ach release in a head-fixed mouse on a wheel, evoking Ach levels that are nearly five-fold higher than those evoked during the 2ABT (**Extended Data Fig. 6a**). CIN activation during the 2ABT in a subset of trials increased DA release in both rewarded and unrewarded trials (**Extended Data Fig. 6b and 6c**); however, the effect is small compared to the magnitudes of CIN-evoked DA observed *in vitro* and of reward-dependent DA release *in vivo*.

We turned to loss of function experiments to assess if Ach release from CINs significantly contributes to DA dynamics during behavior. To abolish neurotransmission in CINs, we used Cre-conditional AAVs and *Chat-IRES-Cre* mice to selectively express tetanus toxin (TelC) and prevent synaptic vesicle fusion in CINs (**Fig. 3a and 3b**). Optogenetic activation of CINs in control striatal slices evokes DA release as measured by carbon fiber amperometry; however, this is abolished if CINs express TelC (**Fig. 3c**). Furthermore, Ach transients during rewarded trials are greatly suppressed by TelC expression in CINs compared to expression of a control protein, mCherry (**Fig. 3d**). Thus, TelC potently inhibits Ach release from CINs *in vitro* and *in vivo*.

**Fig. 3.**
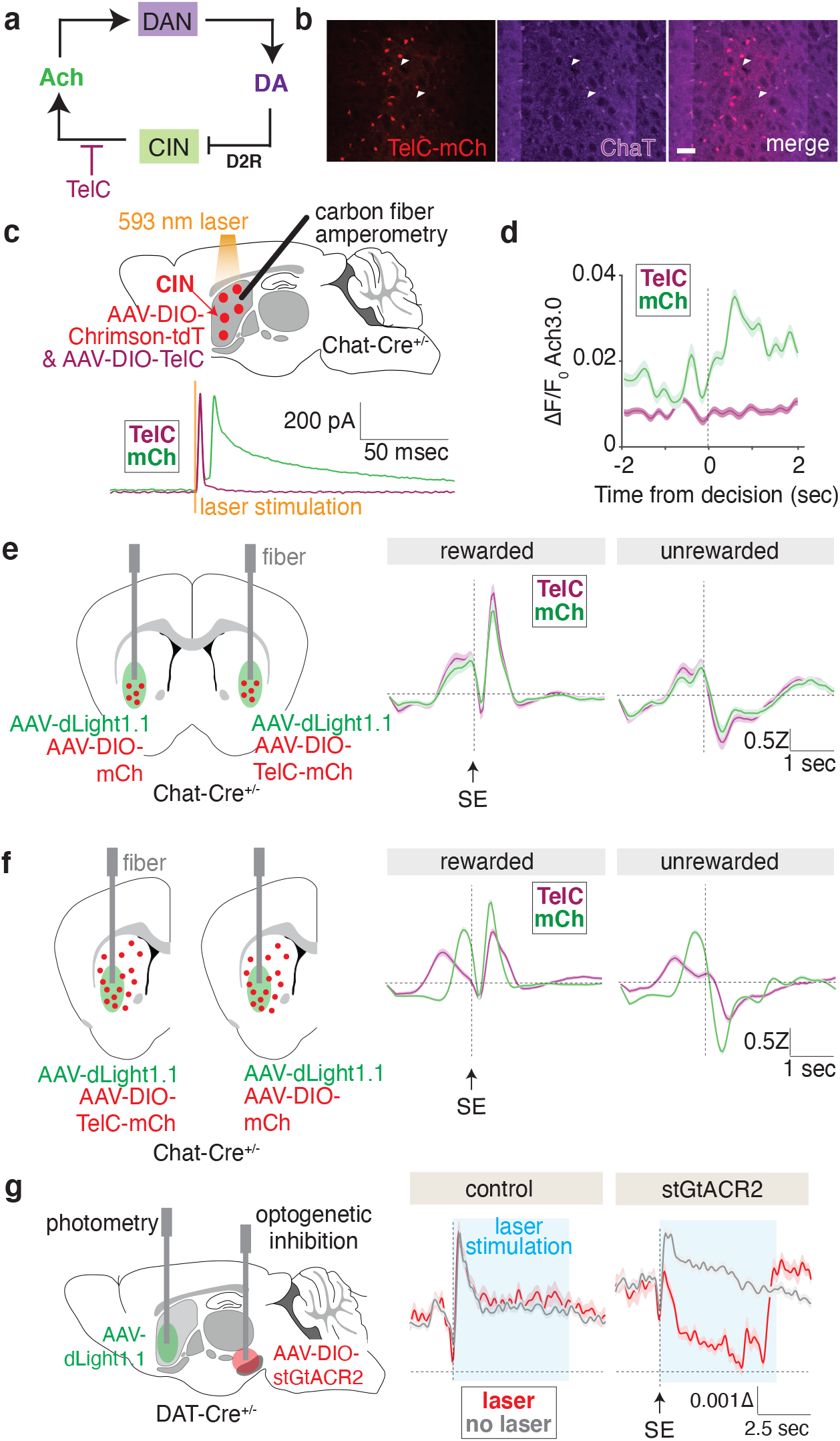
Ach does not regulate DA dynamics during decision-making. **a**. Schematic of the tetanus toxin (TelC) perturbation of Ach release. **b**. Epifluorescence images of tetanus toxin linked to mCherry (TelC-mCh) expressed in choline acetyltransferase (ChaT)-positive cells in the VLS. Scale bar (white): 200 μm. **c**. Carbon fiber amperometry in an acute striatal slice containing CINs coexpressing Chrimson with either a control protein mCherry (mCh) or TelC. Schematic of the experimental setup (top) and amperometry recordings aligned to laser stimulation (bottom). **d**. Ach release in VLS during rewarded trials recorded with fiber photometry in mice with CINs expressing mCh or TelC. The average ΔF/F_0_ of the sensor signal (bold line) is aligned to side port entry (SE) with standard error of the mean (SEM) overlaid (shaded region) (n = 3 mice). **e**. Injections and implantations for fiber photometry experiments to measure DA release in the presence of CIN-specific expression of TelC or mCh in separate hemispheres of the same brain (left). DA release during rewarded and unrewarded trials from these mice (middle and right). The side-entry (SE) aligned average Z score of the sensor signal is shown (bold line) with SEM overlaid (shaded region) (n = 4 mice). **f**. Injections and implantations for fiber photometry to measure DA release in the context of striatum-wide expression of mCh or TelC in CINs (left). Unilateral injections were performed in two separate cohorts of mice. DA release during rewarded and unrewarded trials from these mice (middle and right). Data are depicted as in (**e**) (n = 5 mice per condition). **g**. Optogenetic inhibition of midbrain DAN cell bodies with simultaneous recordings of DA release in VLS. Schematic of the injections and implantations (left), and summary of DA release during the 2ABT from mice lacking opsin expression (middle) or mice expressing the inhibitory opsin stGtACR2 (right). Average ΔF/F_0_ of dlight1.1 is indicated (bold line) with SEM overlaid (shaded region) (n = 3 mice).

In mice performing the 2ABT, we inhibited Ach release in the VLS of one hemisphere using TelC and compared DA release to the VLS within the other hemisphere in which CINs express a control protein (**Fig. 3e, left; Extended Data Fig. 7a**). Surprisingly, loss of Ach release did not perturb DA transients in rewarded or unrewarded trials recorded with dLight1.1 (**Fig. 3e**) or with rDAh in dual recordings in which we validate the suppression of Ach release (**Extended Data Fig. 7c**). In addition to comparing Z scored DA signals, we also analyzed the amplitudes of DA sensor fluorescence transients (ΔF/F_0_) to address if TelC lowers the overall magnitude of DA release throughout the trial. The average fluorescence of dLight1.1 is larger in the hemisphere in which CINs express TelC (**Extended Data Fig. 7b, right**); however, this is difficult to interpret given the high variability of ΔF/F_0_ across individual mice which can stem from varying autofluorescence and sensor expression levels (**Extended Data Fig. 7b, left**). Consistent with the lack of effect of Ach loss on DA dynamics in VLS, mice did not exhibit behavioral deficits except for a slight change in decision time (time from center port entry to side port entry) (**Extended Data Fig. 8a)**. All other performance metrics such as reward rate, lose-switch rate, RFLR model fits, and the dynamics of port switching and high port occupancy were not significantly altered (**Extended Data Fig. 8a – 8d**).

Because the disruption of Ach release was limited to the VLS, the lack of effect on DA release may result from sustained synchronized CIN activity outside the VLS, which may be sufficient to drive DA release within the VLS. To address this possibility, we expressed TelC in CINs throughout a large fraction of the striatum using a multi-site injection approach (**Fig. 3f, left; Extended Data Fig. 7d**). This widespread loss of Ach release affected behaviorally-evoked DA signals (**Fig. 3f**) but also induces severe behavioral defects, including impaired locomotion (**Extended Data Fig. 8e**), impaired switching rates and dynamics (**Extended Data Fig. 8f, 8h**), altered *α* (stickiness in action selection) and *β* (weight given to new information) RFLR coefficients (**Extended Data Fig. 8g**). In addition, all GLM kernels underlying DA dynamics were changed by widespread TelC expression (**Extended Data Fig. 7e**) (mCh DA GLM R^2^ = 0.220; TelC DA GLM R^2^ = 0.075). Although these perturbations confirm the importance of CIN-dependent Ach release for proper performance in this task, they make it difficult to discern if the perturbations of DA signals are due to Ach loss or due to behavioral alterations caused by Ach loss. However, despite severe striatum-wide loss in Ach, RPE encoding is still sustained by DA, such that DA rises during rewarded trials and falls during unrewarded trials (**Fig. 3f**) and the GLM kernels associated with side entry and reward maintain their opposing polarity (**Extended Data Fig. 7e**). Thus, the RPE encoding features of DA do not require CIN-mediated release but rather likely depend on changes in action potential firing generated in the SNc and VTA. Indeed, while suppression of Ach release did not alter DA levels, modulation of DAN activity does so robustly, as evidenced by a significant reduction in DA levels upon optogenetic inhibition of DANs via stGtACR2 (**Fig. 3g**). Altogether, we find that loss of Ach release within the VLS does not impair DA dynamics such that, although modulation of CIN activity is sufficient to drive DA release *in vivo*, the context in which it does so remains to be determined.

### D2Rs are necessary for DA-dependent inhibition of Ach levels

During a trial, there are two periods in which Ach levels are depressed while DA levels rise: first, as the mice move from center to side port, and second, during rewarded trials following side port entry (**Fig. 2b**). Because these opposite signed changes in DA and Ach signals coincide, we hypothesize that D2Rs may mediate the depression of Ach levels. To test this possibility, we assayed if optogenetic manipulations of DA neurons *in vivo* affect striatal Ach levels and if any such effects are D2R-dependent. We increased and decreased DA levels in the VLS via photoactivation of excitatory and inhibitory optogenetic proteins Chrimson and Jaws^56,57^, respectively, expressed in SNc and VTA DANs in a head-fixed mouse on a wheel (**Fig. 4a**). These manipulations altered Ach levels in the direction opposite to optogenetically-evoked changes in DA levels, consistent with DA inhibiting Ach release (**Fig. 4b and 4c, left**). Importantly, these effects are D2R-dependent as they are abolished by administration of eticlopride, a D2R antagonist (**Fig. 4b and 4c, right**). Thus, changes in DA are sufficient to bidirectionally regulate Ach levels *in vivo*, consistent with basal engagement and dynamic modulation of DA-dependent inhibition of CINs.

**Fig. 4.**
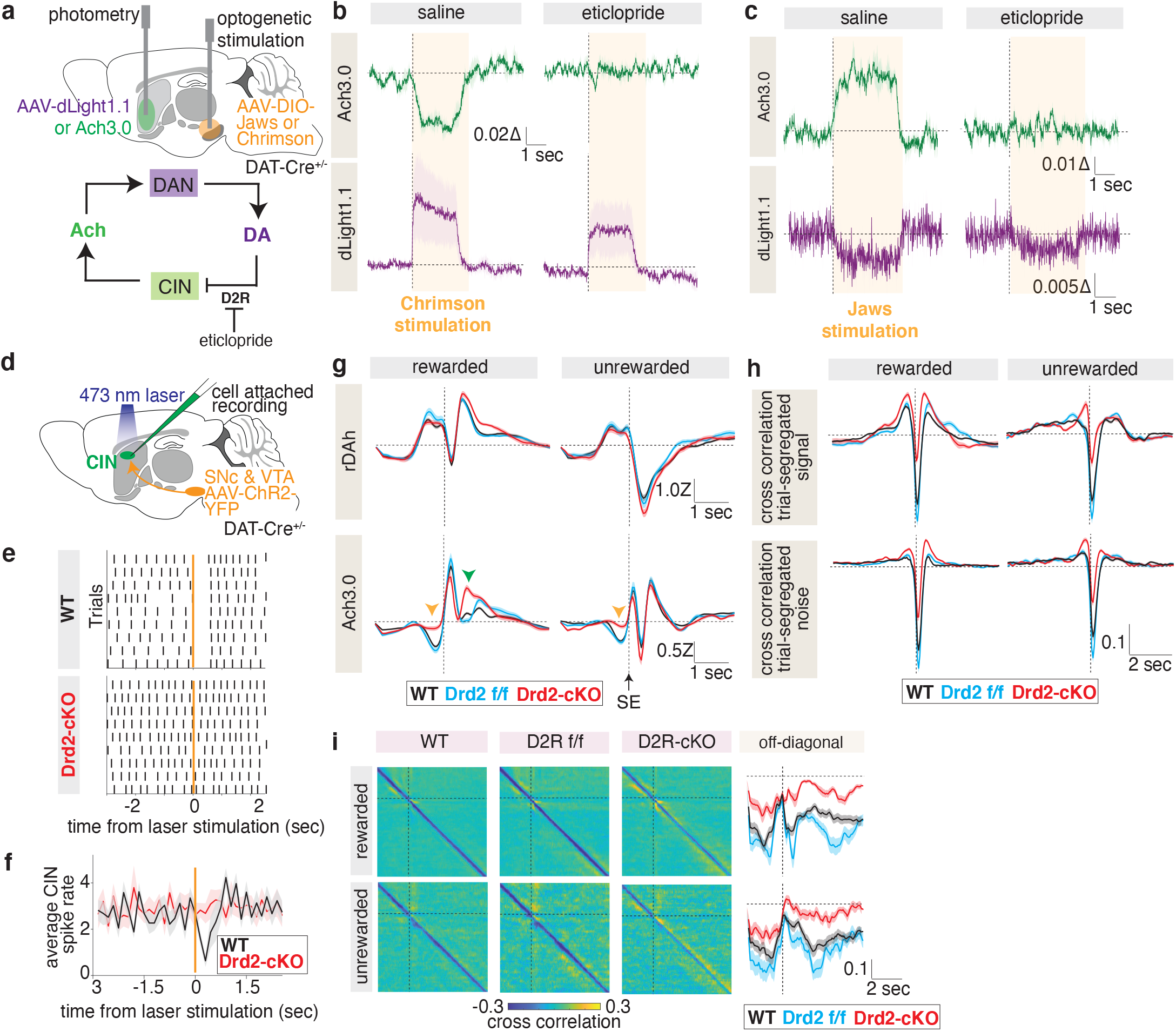
D2Rs are required for DA-mediated inhibition of Ach signals. **a**. Schematic illustrating the injection and fiber implantation strategy (top). Eticlopride, a D2R antagonist, was applied during these recordings (bottom). **b**. Ach and DA release during Chrimson-mediated optogenetic activation of DAN somas with an intraperitoneal injection of saline (left) or eticlopride (right) prior to the recording. Average ΔF/F_0_ is shown (bold lines) with standard error of the mean (SEM) overlaid (shaded region) (n = 3 mice). Data are aligned to the onset of laser stimulation (dashed vertical line). **c**. Ach and DA release during Jaws-mediated optogenetic inhibition of DAN somas with an intraperitoneal injection of saline (left) or eticlopride (right) prior to the recording. Data are depicted as in (**b**) (n = 3 mice). **d**. Schematic illustrating the injection and recording setup to determine the effect of DA release on CIN firing in an acute striatal slice. ChR2-positive DAN afferents (orange) are activated with a 473 nm laser pulse, and CIN firing is recorded with a cell attached pipette (green). **e**. Representative single cell responses to optogenetic activation (orange) of ChR2-positive DAN afferents recorded from CINs with D2R expression (WT) or without it (Drd2-cKO). Each bold vertical line denotes a CIN action potential. **f**. Population perievent spike histograms of average spike discharge (solid lines) with SEM (shaded region) for cells recorded in (**d**) (WT: n = 9 cells; Drd2-cKO: n = 9 cells). **g**. Simultaneous trial-averaged Ach and DA recordings from the indicated genotypes, aligned to side port entry (SE). Average signals are depicted (bold) with SEM overlaid (shaded region) (WT: n = 12 mice; Drd2 f/f: n = 7 mice; Drd2-cKO: n = 8 mice). An orange arrow denotes the increase in Ach signal that occurs when the mice move from center to side port, and a green arrow denotes this change in Ach signal after side port entry. **h**. Covariance of DA and Ach dynamics in mice shown in (**g**), in which DA lags Ach by the indicated time. Analyses of the trial-segregated signals (top) and their residuals (bottom) are shown for the indicated trial types. Average covariance is depicted (bold line) with SEM outlined (shaded region). **i**. Moment-to-moment covariance matrix for mice in (**g**) of DA and Ach release in rewarded and unrewarded trials with their respective off-diagonal signals (bold lines) and SEM overlaid (shaded region) (far right).

Because D2Rs are expressed by other cell types in the brain and D2R-blockade has significant behavioral effects that prevent mice from performing the task^58^, we used an alternative method to determine if DA suppresses Ach release during the task. We employed a genetic strategy to knock-out D2Rs specifically in CINs and refer to this transgenic mouse line (*Chat-IRES-Cre; Drd2*^*f/f*^) as Drd2-cKO.

To confirm functional loss of D2Rs in CINs, we compared the ability of DA to reduce CIN firing in striatal slices from wild type versus Drd2-cKO mice (**Fig. 4d**). In wild type mice, release of DA following laser stimulation of channelrhodopsin-expressing DA neuron terminals robustly reduced CIN firing, as measured by cell attached recordings, but this effect is absent in CINs from Drd2-cKO mice (**Fig. 4e and 4f**).

To determine how D2R loss in CINs affects Ach release during the 2ABT, we compared neuromodulator dynamics in the VLS of Drd2-cKO CIN mice with that in two control groups: *Chat-IRES-Cre* mice (referred to as WT), and *Drd2*-floxed mice (referred to as Drd2-ff). We found that loss of D2Rs in CINs abolished both instances of Ach suppression that coincide with a rise in DA levels (**Fig. 4g, bottom**): the Ach dip that normally occurs in all trials during center to side port transition is lost (**Fig. 4g, orange arrow**), and an additional peak of Ach emerges following side port entry in rewarded trials (**Fig. 4g, green arrow**). Importantly, these changes occurred despite relatively unperturbed DA dynamics during the task across the three genotypes (**Fig. 4g, top**). Modeling these signals with a history GLM recapitulates these effects – DA kernels are comparable across the three cohorts (**Extended Data Fig. 9c**). In contrast, all side-entry Ach kernels are greatly altered in Drd2-cKO mice, exhibiting an increase in *β* coefficients at negative time shifts across all histories as well as an increase in *β* coefficients at positive time shifts for histories with a rewarded outcome on the current trial (**Extended Data Fig. 9b**) (WT DA GLM R^2^ = 0.129; Drd2-ff DA GLM R^2^ = 0.122, Drd2-cKO DA GLM R^2^ = 0.155; WT Ach GLM R^2^ = 0.202; Drd2-ff Ach GLM R^2^ = 0.262, Drd2-cKO Ach GLM R^2^ = 0.214). As a result, the relationship between Ach and DA signals in Drd2-cKO mice is severely disrupted (**Extended Data Fig. 9a**), with a significant reduction in both the magnitude of their anticorrelation (**Fig. 4h**) and the off-diagonal negative covariance (**Fig. 4i**). Thus, D2Rs are required for DA to inhibit Ach release *in vivo*.

To determine if D2R loss in CINs affects decision-making, we assessed the performance of Drd2-cKO mice in the 2ABT. General performance metrics (**Extended Data Fig. 10a**), block transition dynamics (**Extended Data Fig. S10b and S10c**), and RFLR coefficients (**Extended Data Fig. 10d**) are comparable across the three genotypes. Although Drd2-cKO mice exhibit a slightly altered decision time, lose-switch rate, and *α* and *β* RFLR coefficients compared to Drd2-ff mice, this is not observed with WT mice, suggesting that there are baseline behavioral differences between the two control groups. However, consistent differences emerge between Drd2-cKO mice and both control groups when performance is parsed by history. Drd2-cKO mice are impaired in their ability to switch across multiple histories when compared to both Drd2-ff and WT cohorts, with significantly reduced switching rates in ab and Aa sequences (**Extended Data Fig. 10e**). This reveals a previously unknown role for D2R-dependent Ach pauses in enabling past choices and outcomes to promote complex changes in behavior. In conclusion, we find that D2Rs are required for DA to repress Ach levels during precise moments within a trial, and loss of this regulation impairs the ability of mice to modify switching behavior.

### Cortical and thalamic inputs drive changes in striatal Ach levels

Although DA shapes Ach signals during decision-making, we observe additional fluctuations in Ach that are DA-independent. For example, in unrewarded trials, Ach signals remain repressed following side port entry, even in mice in which CINs lack D2Rs (**Fig. 4g, lower right**). Furthermore, additional inputs are required to drive the increases in Ach that occur upon side port entry and during the consumption period. Finally, the momentary disruption of the negative covariance between Ach and DA signals points to the existence of other factors that can alter Ach and DA dynamics (**Fig. 2e and 2f**).

To discover other potential sources of regulation of striatal Ach, we examined inputs to striatum from the cortex and thalamus, both of which synapse onto CINs and modulate their firing rates^43,44,59^. To determine if these regions project into the VLS, we performed retrograde tracing with cholera toxin. We find that a broad distribution of cells from multiple cortical regions send afferents into the VLS (**Fig. 5a**). Meanwhile, thalamic inputs into this striatal region originate predominantly from the parafascicular nucleus.

**Fig. 5.**
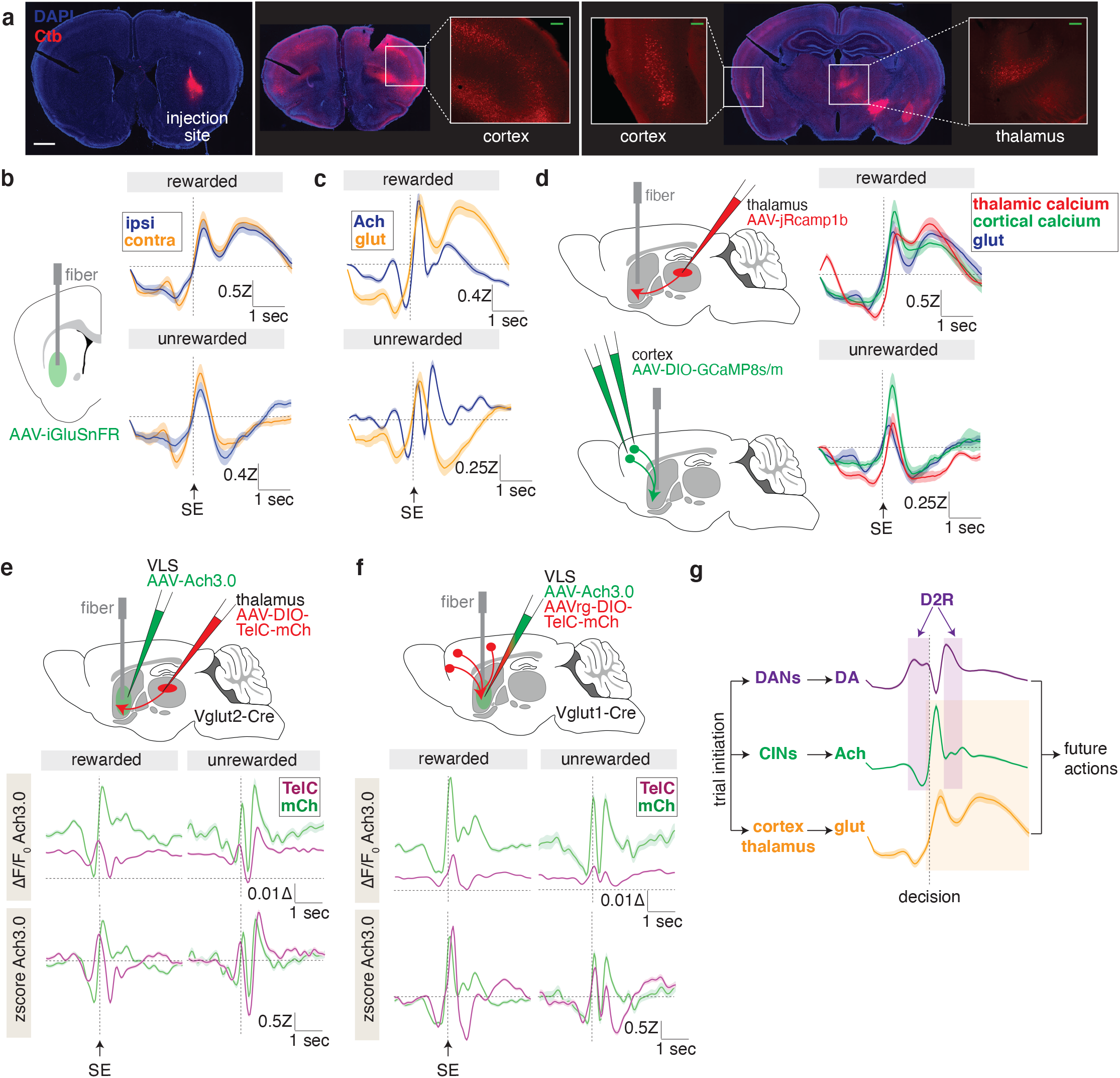
Corticostriatal and thalamostriatal inputs are necessary to drive Ach signals in the VLS. **a**. Retrograde labelling of cortex and thalamus with cholera toxin (Ctb) injected in the VLS. A representative mouse is shown. Epifluorescence images of the injection site (left) and representative coronal sections that depict Ctb-positive cortical cell bodies (middle) and thalamic cell bodies (right). Scale bar (white): 1 mm; Scale bar (green): 200 μm. **b**. Glutamate release from ipsilateral versus contralateral 2ABT trials measured with the iGluSnFR sensor (left). Average signals (bold lines) are aligned to side port entry (SE) with standard error of the mean (SEM) overlaid (shaded region) (right) (n = 4 mice). **b**. Overlay of glutamate and Ach release recorded from separate mice during the indicated trial types. Data is presented as in (**b**) (glutamate: n = 4 mice; Ach: n = 9 mice). **c**. Injection and fiber implantation scheme for photometry of calcium dynamics in thalamic (upper left) and cortical (lower left) terminals in the VLS. Two injections were required to cover the medial-lateral spread of cortical cells that project to VLS. Overlay of thalamic calcium, cortical calcium, and VLS glutamate signals (right) in the VLS of separate mice for the indicated trial types. Data are depicted as in (**b**) (glutamate: n = 4 mice; thalamic calcium: n = 6 mice; cortical calcium: n=5 mice). **d**. Injection strategy and fiber implantation for tetanus (TelC) or mCherry (mCh) expression in thalamus with Ach recordings in VLS (top). ΔF/F_0_ (middle) and Z scored signals (bottom) of the mean Ach release from the indicated treatment groups (TelC: tetanus; mCh: mCherry). Data are depicted as in (**b**) (mCh: n = 3 mice; TelC: n = 5 mice). **e**. Injection strategy and fiber implantation for TelC or mCh expression in cortex with Ach recordings in VLS (top). ΔF/F_0_ (middle) and Z scored signals (bottom) of the average Ach release from the indicated treatment groups. Data are depicted as in (**b**) (mCh: n = 4 mice; TelC: n = 5 mice). **f**. Summary of findings. Trial initiation evokes the release of multiple neurotransmitters in the VLS, all of which interact and influence decision making. Glutamate release from cortex and thalamus are necessary to promote Ach release (orange shaded box), while DA release inhibits it at specific trial moments via D2Rs (purple shaded boxes). Altogether, this guides future actions.

To assess the potential influence of glutamatergic inputs from cortex and thalamus on striatal Ach, we first determined if striatal glutamate and Ach signals are correlated. Using the glutamate sensor iGluSnFR^60^, we find that striatal glutamate levels vary during the task but are not lateralized (**Fig. 5b**), allowing us to again combine signals from ipsiversive and contraversive trials. Glutamate signals are suppressed prior to the choice and rise during side port entry in an analogous manner to Ach (**Fig. 5c**). In rewarded trials, glutamate exhibits an additional phase of sustained increase during consumption, which is absent in unrewarded trials (**Fig. 5c**).

The activities of cortical and thalamic terminals in VLS, measured with genetically encoded calcium sensors, coincide with glutamate release across both trial types, suggesting that both inputs can contribute to glutamate release in this region (**Fig. 5d, right**). To detect thalamic activity, we expressed jRCaMP1b, a red calcium indicator, in thalamic somas of separate animals and recorded terminal activity in VLS (**Fig. 5d, upper left**). Timing offsets between jRCaMP1b and Ach3.0 signals are likely due to varying sensor kinetics ^53,61^. Meanwhile, a multisite injection strategy was required for cortical inputs given their widespread distribution (**Fig. 5d, bottom left**). In addition, we found that a brighter calcium sensor, GCaMP8, was necessary for detection of cortical signals arising from these dispersed sources. Altogether, these data demonstrate substantial coincident dynamics from glutamatergic inputs into striatum, providing a basis for the possibility that cortical and thalamic inputs can drive changes in Ach levels.

To test whether each input is required for Ach release, we expressed TelC unilaterally in the thalamus or cortex. For cortical inputs, we used a retrograde AAV approach in *Vglut1-IRES-Cre* mice to restrict expression of the toxin in only cells that project into VLS (**Fig. 5f top, Extended Data Fig. 11b**). However, for thalamic inputs, we could not use a retrograde approach in *Vglut2-IRES-Cre* mice due to Vglut2 expression in cortical regions (Allen Institute); therefore, we instead injected TelC directly into the thalamus (**Fig. 5e top, Extended Data Fig. 11a**). This perturbation impaired the performance of animals on the 2ABT, reflecting the importance of both inputs in regulating striatal function. Although TelC-expressing animals could achieve a comparable, albeit lower, reward rate than control mice, they exhibited impaired switch dynamics following rewarded trials (**Extended Data Fig. 12a and 12e**), block transitions (**Extended Data Fig. 12b and 12f**) and across multiple choice-outcome histories (**Extended Data Fig. 12c and 12g**). Consistent with this impairment, the RFLR model description revealed a reduction in the *β* and an increase in *τ* reflecting weakened incorporation of new evidence and faster information decay, respectively (**Extended Data Fig. 12d and 12h**). Mice with thalamic TelC expression performed more poorly than those with cortical manipulations. This may be because a large majority of thalamic inputs into striatum received TelC whereas for cortical inputs, only those that project into VLS expressed the toxin.

Loss of neurotransmission from each region robustly dampened Ach transients across all trials, as seen in the lowered ΔF/F_0_ of the Ach sensor signal (**Fig. 5e and 5f, bottom**). The degree of suppression of ΔF/F_0_ was strong and consistent across mice and sufficient to overcome any underlying variability in signals, unlike CIN perturbation of DA levels (**Extended Data Fig. 11c**). Analysis of the remaining Ach transients reveals unique ways that cortical and thalamic inputs modulate Ach levels. In unrewarded trials, loss of cortical but not thalamic inputs perturbs Ach transients following side port entry, suggesting a specific role for the cortex in driving this signal. In addition, only loss of the thalamic input shifts the timing of Ach release, which may reflect the greater degree of behavioral disruption in mice with thalamic TelC injections. These results point to cortical and thalamic inputs as candidates for driving the momentary disruption in covariance between Ach and DA (**Fig. 2e**). Testing this hypothesis requires perturbation of these inputs; however, this results in altered Ach signals, which in turn prevents calculation of the covariance between DA and Ach. Overall, our results reveal that both the cortex and thalamus are required to sustain Ach levels during decision-making, and each input can uniquely alter the dynamics of striatal Ach during a trial.

## Discussion

DA and Ach are critical neurotransmitters that directly modulate each other’s release *in vitro*. However, whether these interactions regulate neurotransmitter levels *in vivo*, particularly during decision making, is not known. To address this, we evaluated how striatal DA and Ach dynamics are regulated by the proposed bidirectional circuit during a task that requires mice to make choices flexibly within a changing environment. We revealed that DA and Ach signals are generally anticorrelated across time, but that this relationship is dynamic and modulated by action, reward, and their histories. Whereas striatal Ach release does not modulate DA dynamics, DA exerts a key influence on Ach signals via D2Rs. Without this interaction, the ability of action and reward history to influence decision-making is diminished. In addition to DA, cortical and thalamic inputs concurrently drive Ach release throughout the trial. Altogether, we define precise roles for local and long-range glutamatergic inputs in modulating striatal Ach signals (**Fig. 5g**), and we highlight the complex coordination of multiple neurotransmitter interactions during decision-making.

### CIN regulation of striatal DA

Although synchronous CIN activation robustly triggers DA release *in vitro*, we did not find evidence that this effect is functionally important *in vivo* in the behavioral context of a reward-guided behavior: DA dynamics are sustained despite loss of Ach in the VLS, and RPE encoding remains even in the absence of Ach release throughout the entire striatum. Nevertheless, specific deletion of muscarinic and nicotinic receptors in DA neurons are necessary to reveal if Ach can regulate other aspects of DA release. Furthermore, we found that optogenetic activation of CINs, despite being able to drive large transients in Ach levels, caused only small changes DA levels *in vivo* (e.g., compared to those evoked by rewards), which contrasts with the robust DA release evoked in striatal slices^32,62^. One caveat is that our method may not synchronously activate a sufficient population of CINs due to the spatial constraints of opsin expression and laser excitation. The discrepancy between the ability of CINs to regulate DA release *in vitro* and *in vivo* is surprising and may stem from fundamental differences between the two experimental systems. Within a striatal slice, basal neuromodulator levels are low, whereas *in vivo* CINs and DANs are both spontaneously active and constantly modulate their activity in response to environmental cues and stimuli, which may create less permissive conditions for CINs to effectively drive DA release. Furthermore, *in vivo* CINs may be more inhibited via D2Rs, nAChRs may be more desensitized, or CIN activity may not be sufficiently synchronized across striatum. However, it remains possible that CINs can modulate DA levels to a greater extent in other *in vivo* contexts, for instance, set shifting^63^ and reversal and extinction learning^64^.

### DA regulation of striatal Ach

CIN pauses emerge after classical conditioning in response to salient and reward-predicting cues^24^, and their coincidence with and dependence on DA release support the requirement for DA to generate this pause. However, a long-standing debate remains about whether DA is responsible for CIN pauses *in vivo*. Subsequent studies found that changes in DA firing do not coincide with changes in CIN firing^25^ and Ach pauses were partially repressed, but not eliminated, upon D2R deletion in CINs^46^. Other inputs have been implicated in generating these pauses, including withdrawal of cortical inputs, and excitation of thalamic neurons and GABAergic neurons from the VTA^43,44,65^.

In the 2ABT, we observe Ach dynamics that are consistent with the classical CIN pause. Importantly, we find that Ach signals are reward responsive and sensitive to RPE, which contrasts with previous studies. This difference could stem from the fact that we measured neurotransmitter levels instead of cell firing, and the former may be a more sensitive measure of changes induced by reward outcome. During a trial, we find that a subset of reductions in Ach transients cooccur with increases in DA levels and, contrary to prior findings, are eliminated when D2Rs are deleted in CINs. However, not all reductions in Ach signals are DA-dependent – only those that coincide with a rise in DA release are. In fact, during unrewarded trials Ach levels dip alongside DA following side port entry and this transient is unaffected by D2R loss. Taken together, we posit that not all transient reductions in Ach levels are equivalent, and multiple mechanisms can generate them in a context-dependent manner. Some pauses are DA-dependent, such as those in rewarded trials of the 2ABT, while others may partially depend on DA^46^ or be DA-independent. In future studies, performing these D2R-deletions selectively in CINs in adulthood using a CRISPR-based approach will complement these results and address concerns of circuit rewiring or compensation.

### Extra-striatal regulation of cholinergic Ach

Another long-standing debate has been the role of cortex versus thalamus on CIN activity. Some studies support the thalamus as being the dominant CIN glutamatergic input. Optogenetic studies revealed strong thalamic but negligible cortical connectivity onto CINs^66^, and cortical activity did not correlate with striatal CIN activity in large scale recordings of mice performing a visually guided task^67^. In contrast, other studies have found that both the cortex and thalamus can drive CIN activity *in vitro*^68^ and *in vivo*^59^, and a rabies-based anatomical study revealed that CINs receive extensive inputs from both regions^69^. To address these divergent findings, we silenced each input individually using tetanus toxin. We found that loss of either cortical or thalamic transmission results in severe suppression of Ach levels and disruption of its release patterns; thus, both inputs are important modulators of CINs in the VLS. With our strategy, we cannot determine the relative contribution of each input to CIN activity because the degree of tetanus inhibition may vary due to differences in viral infectivity and toxin efficacy across cell types. Furthermore, disruption of each input affects many cells in the striatum (and in other brain regions) such that some of the effects on Ach levels may be indirect. Finally, whether both inputs play a similar role in other striatal regions and in other behavioral contexts remains to be determined. Given the spatial and functional heterogeneity of striatum, a comprehensive survey of these inputs is necessary to resolve different conclusions across past studies.

### Cholinergic contributions to behavior

From our study, we gained a snapshot of only a few of the multitude of complex interactions that take place in striatum. Much more remains to be understood about how neurotransmitter release is integrated across time and space to direct striatal function. It has been hypothesized that CIN pauses are time windows that allow DA to potentiate corticostriatal and thalamostriatal synapses^70^. Consistent with this, a precise *in vivo* coincidence of CIN pauses, DAN activation, and striatal spiny projection neuron (SPN) depolarization is required for long-term potentiation of corticostriatal synapses with SPNs^71^. In the 2ABT, different reward outcomes yield unique patterns of neurotransmitter release, the combination of which could tune the plasticity of striatal synapses in distinct ways to drive behavior. During rewarded trials, DA and Ach are tightly anticorrelated, which could permit potentiation at multiple time points. In unrewarded trials, DA and Ach release are repressed, which could in turn promote depression. With D2R loss, the temporal gating of DA by Ach is disrupted, resulting in aberrant Ach signaling that may inhibit plasticity by reducing glutamatergic transmission^72^ and ultimately impair the ability of mice to incorporate past knowledge into current actions. In conclusion, by using a diverse toolset to interrogate and alter neurotransmitter levels during a complex behavioral task, we establish a more precise *in vivo* role for a long-defined *in vitro* circuit and reveal new modes of CIN regulation by local and long-range inputs. Moreover, our findings set the framework for further studies upon which we can gain a deeper understanding of the neurochemical basis of decision making and behavior.

## Methods

### Mice

The following mice lines were used: C57BL6/J (Jackson labs #000664); *ChAT-IRES-Cre* (Jackson labs #006410); *DAT-IRES-Cre* (Jackson labs #006660), *Drd2*^*loxP*^ (Jackson labs #020631); *Vglut2-IRES-Cre* (Jackson labs #028863); *Vglut1-IRES-Cre* (Jackson labs #023527). All mice were bred on a C57BL/6J genetic background and heterozygotes were used unless noted. For behavior experiments, males at 6-8 weeks of age were used. Only males were used to avoid any behavioral variation due to the estrous cycle in female mice and because of recent findings that only male behavior is affected by loss of muscarinic Ach receptors^73^. All animal care and experimental manipulations were performed in accordance with protocols approved by the Harvard Standing Committee on Animal Care, following guidelines described in the US NIH Guide for the Care and Use of Laboratory Animals.

### Intracranial injections

Mice were anesthetized with 5% isoflurane and maintained under surgery with 1.5% isoflurane and 0.08% O_2_. Under the stereotaxic frame (David Kopf Instruments), the skull was exposed in aseptic conditions, a small craniotomy (∼300 um) was drilled, and the virus was injected into the following regions with the associated coordinates listed from bregma: VLS (coordinates: 0.6 mm A/P, -/+2.3 mm M/L, and 3.2 mm D/V); SNc and VTA (coordinates: -3.35 mm A/P, -/+ 1.75 mm M/L, and 4.3 mm D/V); thalamus (coordinates: -2.1 mm A/P, -/+1.0 mm M/L, and 3.5 mm D/V); prefrontal cortex (coordinates: 2.0 mm A/P, -/+0.4 mm M/L, and 2.3 mm D/V).

Injections were performed as previously described^74^. A pulled glass pipette was held in the brain for 3 minutes, and viruses were infused at a rate of 50 nl/min (VLS), 30-40 nl/min (PFC), and 70 nl/min (SNc/VTA) with a syringe pump (Harvard Apparatus, #883015). Pipettes were slowly withdrawn (<10 μm/s) at least 6 min after the end of the infusion. 350 nl was infused per injection site except for Ctb 555 injections (50 nl at 4 μg/μl).

For AAV injections, the wound was sutured. For fiber implants, following AAV injection, the skull was scored lightly with a razor blade to promote glue adhesion. Then, a 200 μm blunt end fiber (MFC_200/230-0.48_4 mm, Doric Lenses) was slowly inserted into the brain until it reached 100 μm above injection site. The fiber was held in place with glue (Loctite gel #454) and hardening accelerated with application of Zip Kicker (Pacer Technology). A metal headplate was glued at lambda and white cement (Parkell) was applied on top of the glue to further secure the headplate and fibers. Fiber implants were protected with a removable plastic cap (Doric Lenses) until recordings.

Following the surgery, mice were placed in a cage with a heating pad until their activity was recovered before returning to their home cage. Mice were given pre- and post-operative oral carprofen (CPF, 5 mg/kg/day) as an analgesic and monitored daily for at least 4 days post-surgery. At least 4 weeks passed after virus injection before experiments were performed, except for retrograde tracer injections in which 1 week passed.

### Viruses

The following viruses were used, with source and titer indicated in parentheses:

AAV2/9 hSyn-dlight1.1 (Boston’s Children’s hospital core (BCH); 6E12 GC/ml)

AAV2/9 hSyn-dlight1.1 D103A (BCH, 2.5E12 GC/ml)

AAV2/9 hSyn-GRAB Ach3.0 (WZ Biosciences, 6.5E12 GC/ml)

AAV2/9 hSyn-GRAB Ach3.0 mutant (Vigene, 1E12 GC/ml)

AAV 2/9 hSyn-GRAB-rDA1h (WZ Biosciences, 1.9E13 GC/ml)

AAV2/1 hSyn-DIO-ChrimsonR-tdTomato (UNC Vector Core, 2E12 GC/ml)

AAV2/8 hSyn-SIO-TelC-mCherry (Janelia Viral Core, 2.2E12 GC/ml)

AAV2/rg hSyn-SIO-TelC-mCherry (Janelia Viral Core, 5.5E12 GC/ml)

AAV2/8 hSyn-DIO-mCherry (Addgene, 2.5E13 GC/ml)

AAV2/rg hSyn-DIO-mCherry (Addgene, 9E12 GC/ml)

AAV2/1 hSyn-DIO-stGtACR2-FusionRed (Addgene, 4.2E13 GC/ml)

AAV2/1 hSyn-SF-iGluSnFR.A1848 (Addgene, 3E12 GC/ml)

AAV2/1 hSyn-DIO-NES-jRcamp1b-WPRE-SV40 (Addgene, 4.5E13 GC/ml)

AAV2/1 hsyn-jGCaMP8s-WPRE (Addgene, 2.8E13 GC/ml)

AAV2/1 hsyn-jGCaMP8m-WPRE (Addgene, 2.0E13 GC/ml)

### Immunohistochemistry

Mice were anaesthetized by isoflurane inhalation and transcardially perfused with phosphate-buffered saline (PBS) followed by 4% paraformaldehyde (PFA) in PBS. Brains were extracted and stored in 4% PFA PBS for at least 8 hours or in 4% PFA, 0.02% sodium azide, and PBS for long term storage at 4°C. Brains were sliced into 70 μm thick free-floating sections with a Leica VT1000s vibratome. Selected slices were transferred to a six well plate and rinsed three times for 5 minutes each in PBS. They were then blocked with rotation at room temperature for an hour in blocking buffer (5% normal goat serum (Abcam), 0.2% TritonX-100 PBS). Blocking buffer was removed and replaced with 500-700 μl of a solution containing the indicated primary antibody. Slices were incubated overnight with side-to-side rotation at 4°C. The next day, slices were transferred to a clean well and washed five times, 5 minutes each in PBST (PBS with 0.2% TritonX-100). Following the final wash, slices were incubated for 1.5 hours in 500-700 μl of the indicated secondary antibody diluted 1:500 in blocking buffer. Slices were washed four times in PBST for 5 minutes each, then four times in PBS for 5 minutes each before mounting with ProLong Diamond Antifade Mountant with DAPI (Thermo Fisher Scientific). Slices were imaged with an Olympus VS120 slide scanning microscope.

### Primary antibodies

The following antibodies were used with the source and dilution indicated in parentheses:

goat anti-Choline acetyltransferase (Millipore Sigma #AB144P; 1:200)

mouse anti-tyrosine hydroxylase (Immunostar #22941; 1:1000)

chicken anti-GFP (Abcam ab13970; 1:1500)

rabbit anti-GFP (Novus Biologicals #NB600-308; 1:1000)

rabbit anti-mCherry (Takara Bio #632496; 1:1000)

rabbit anti-GFAP (Abcam ab7260; 1:1500)

### Behavior apparatus, training, and task

The apparatus used for the behavior is as described previously^49^ with the following modifications. Clear acrylic barriers 5.5 cm in length were installed in between the center and side ports prior to training to extend the trial time to aid in better resolved photometry recordings. Water was delivered in 3 μl increments. Hardware and software to control the behavior box is available online: https://github.com/HMS-RIC/TwoArmedBandit

Mice were water restricted 1 ml per day prior to training and maintained at ≥80% initial body weight for the full duration of training and photometry. All training sessions were conducted in the dark under red light conditions. A blue LED above the center port signals to the mouse to initiate a trial by poking in the center port. Blue LEDs above the side ports are then activated, signaling the mouse to poke in the left or right side port within 5 seconds. At any given instance, only one side port rewards water. Reward probabilities are defined by custom software (MATLAB). Withdrawal from the side port ends the trial and begins a 1 second intertrial interval (ITI). An expert mouse can perform 200-300 trials in a session.

To train the mice to proficiency, they were subjected to incremental training stages. Each training session lasts for 30-60 minutes, adjusted according to the mouse’s performance. Mice progress to the next stage once they were able to complete at least 100 successful trials with at least a 75% reward rate. On the first day, they were habituated to the behavior box, with water being delivered from both side ports and triggered only by a side port poke. In the next stage, mice learned the trial structure – only a poke in center port followed by a side port poke delivers water. Then, the mice transitioned to learning the block structure, in which 30 successful trials on one side port triggers a deterministically rewarded port (P_high_ = 100%) to switch to the other side port. Finally, mice performed trials in the presence of barriers in between the center and side ports. A series of transparent barriers of increasing size (extra-small (1.5 cm), small (3 cm), medium (4 cm), and long (5.5 cm)) aided in learning. Finally, the mice were trained on probabilistic reward delivery (P_high_ = 95%) and subjected to fiber implantation.

Following fiber implant surgeries, mice were retrained to achieve the same pre-surgery performance level. Habituation to head-fixation on a wheel followed by attachment of a mock photometry patchcord was performed over successive days. Recordings were performed 4 weeks after surgery to allow for stable viral expression levels as well as a consistent and proficient level of task performance from the mice.

### Photometry and behavior recordings

Fiber implants on the mice were connected to a 0.48 NA patchcord (Doric Lenses, MFP_200/220/900-0.48_2m_FCM-MF1.25, low autofluorescence epoxy), which received excitation light and propagated its emission light to a Dorics filter cube (blue excitation light (465-480 nm); red excitation light (555-570 nm), green emission light (500-540 nm); red emission light (580-680 nm) (FMC5_E1(465-480)_F1(500-540) _E2(555-570)_F2(580-680)_S, Doric Lenses). Excitation light originated from LED drivers (Thorlabs) and was amplitude modulated at 167 Hz (470 nm excitation light, M470F3, Thorlabs; LED driver LEDD1B, Thorlabs) and 223 Hz (565 nm excitation light, M565F3, Thorlabs, LED driver LEDD1B, Thorlabs) using MATLAB. The following excitation light powers were used for the indicated sensors: dlight1.1 (25 μW); Ach3.0 (25 μW); rDAh (45 μW); iGluSNFr (15 μW). Signals from the photodetectors were amplified in DC mode with Newport photodetectors or Dorics amplifiers and received by a Labjack (T7) streaming at 2000 Hz. The Labjack also received synchronous information about behavior events logged from the Arduino which controls the behavior box. The following events were recorded: center port entry and exit, side port entry and exit, lick onset and offset, and LED light onset and offset.

### Optogenetic manipulations

For optogenetic stimulations with Chrimson during behavior (**Extended Data Fig. 6c**), 15 mW of 590 nm laser (Optoengine) was evoked in 25% of trials interleaved throughout the session. The excitation light was delivered via the Doric filter cube, which led to a laser stimulation artifact which is removed in the recordings. Only one hemisphere was illuminated in each session. For optogenetic stimulations with Chrimson and with Jaws for DANs (**Fig. 4b and 4c**), 15 mW of 590 nm laser was used, while 0.7 mW of 463 nm laser was used for stGtACR2 stimulations (**Fig. 3g**). For optogenetic stimulations of head-fixed mice on a wheel, in each session, laser excitation duration was 1.5 seconds, with a 45 second intertrial interval, repeated 20 times. Signals displayed are averages of each session (**Fig. 4b and 4c, Extended Data Fig. 6a**). The photometry signal baseline was calculated by averaging the signal 1.5 seconds prior to laser stimulation across the 20 sweeps.

### Behavior performance analysis

Switching behavior of mice in this task is accurately captured by a recursively formulated logistic regression model (RFLR), which depends on an influence of choice history bias (*α*), evidence accumulation (*β*), and the rate of decay of action-outcome information for each trial (*τ*) to make predictions about future choice^49^. Based on our calculated *τ*, a history length of two is sufficient to capture ∼75% of the evidence weight (**Extended Data Fig. 1f**). While a history length of three would increase this evidence weight by 10%, it does so at the cost of reducing the frequency for each of the resulting sequences and limiting our power for estimation. This supports our use of conditional switch probabilities with a history length of two trials for analyses of model performance and behavioral perturbations.

Because we do not know the underlying distribution of our behavioral data, we developed a non-parametric test to determine statistical significance between the experimental and control datasets. We bootstrapped from our original datasets by resampling 1000 times. After each resampling iteration, we calculated the difference (Δ) between the experimental and control groups of the parameter of interest (i.e. RFLR coefficients, reward rate, etc.). This generated a distribution of possible Δs, from which we calculated a 95% confidence interval. If the confidence interval of the Δs did not overlap with zero, which is our null hypothesis, it was annotated as significant.

### Analysis of photometry data

#### Signal demodulation

The frequency modulated signals were detrended using a rolling Z-score with a time window of 1 minute (12000 samples). As the ligand-dependent changes in fluorescence measured *in vivo* are small (few %) and the frequency modulation is large (∼100%), the variance in the frequency modulated signal is largely ligand independent. In addition, the trial structure is rapid with inter-trial intervals of < 3 sec. Thus, Z-scoring on a large time window eliminates photobleaching without affecting signal. Detrended, frequency modulated signals were frequency demodulated by calculating a spectrogram with 1 Hz steps centered on the signal carrier frequency using the MATLAB ‘spectrogram’ function. The spectrogram was calculated in windows of 216 samples with 108 sample overlap, corresponding to a final sampling period of 54 ms. The demodulated signal was calculated as the power averaged across an 8 Hz frequency band centered on the carrier frequency. No additional low-pass filtering was used beyond that introduced by the spectrogram windowing. For quantification of fluorescence transients as Z-scores, the demodulated signal was passed through an additional rolling Z-score (1 min window). In select analyses, (cross-correlation in **Fig. 2d**), the same approach was used but with 72 sample windows with 36 sample overlap in the spectrogram to yield a 18 ms final data sample period.

To synchronize photometry recordings with behavior data, center port entry timestamps from the Arduino were aligned with the digital data stream indicating times of center-port entries. Based on this alignment, all other port and lick timings were aligned and used to calculate the trial-type averaged data shown in all figures. The Z-scored fluorescence signals were averaged across trials, sessions, and mice with no additional data normalization.

#### Cross-variance analyses

Cross-correlation analysis was performed in two ways. First, for cross-correlations with the assumption of semi-wide signals (i.e. as a function of a single variable representing the time shift between two signals), the ‘xcorr’ function in Matlab was used on normalized data and with the “normalized” option set to yield cross-covariance values between -1 and 1, indicating perfect anti-correlation and correlation, respectively. For the continuous demodulated fluorescence, signals were used as input. Shuffling was accomplished by cross-correlating signals across sessions which were truncated to the duration of the shortest signal. For cross-correlation with segregation by trial types, 40 data points before and 60 data points after the event of interest (e.g., center port entry time in all of the cross-correlations shown) were concatenated for all trials of interest (e.g. rewarded trials) in one session and normalized using the xcorr function in MATLAB as above. For noise correlations, the trials-average signal was subtracted trial-by-trial and the residuals were concatenated and treated as above.

Second, for calculation of the two-dimensional cross-covariance (i.e. with no semi-wide assumption), the residual signals following subtraction of the trial-averaged signals were used to calculate the following:

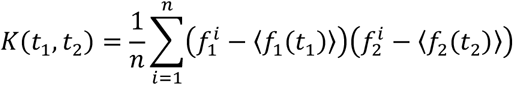

with *i* indicating the trial number (from 1 to n) and *f*_1_ and *f*_2_ corresponding to the two signals analyzed. This corresponds to, for each value of *t*_1_ and *t*_2_, calculating the mean value of the product of the residuals of each signal (relative to its trial-averaged mean) at the corresponding time points.

### General linear model (GLM)

Photometry recordings and behavioral data used for GLMs were collected from the indicated number of mice (see figure legends), with 3-6 sessions per mouse and 150-300 trials per session, of which typically >75% are rewarded. This data was aligned to behavioral events (see *Signal demodulation*) and combined into two time-aligned matrices: a predictive matrix X ∈ R^N x F^, and an associated response vector, y ∈ R^N^, where N represents the number of timesteps recorded in the session and F represents the number of predictive variables used in the analysis. For GLM analyses that included behavioral predictive variables but excluded photometry signals in the predictive matrix, X ∈ {0, 1}^N x F^ as each entry simply indicated if a behavioral event (e.g., a lick) occurred in the time bin.

For each predictive matrix, a design matrix φ(X) ∈ R^N x F(2T+1)^ was constructed to include T time shifts forward and backward (T = 20, 54 ms each), such that the fitting of a standard GLM generated a set of coefficients that comprise the time-dependent kernels for each of the predictive features in X. Some of these time shifts are absent in the first and last T timesteps of each session; thus, those associated 2T data points were excluded, and N was reduced by 2T. To avoid truncation of the center port entry and side port exit kernels, trial boundaries were redefined as T timesteps prior to the center port entry and T timesteps after side port exit. Thus, only data starting shortly before the trial start and after the trial end were modeled (i.e., excluding data from the intertrial interval (ITI) period in which there are no-task relevant behavioral events). For trials where the initial and final time shift spanned the boundary between two trials, the overlapped data was duplicated and included in both trials on either side of the boundary to ensure sufficient representation in training, validation, and test datasets. Because of this and the variability in the ITIs, between ∼1.5% and ∼17.3% of the datapoints used to fit and evaluate the models were present in both the training and test datasets. Finally, the design matrices and response vectors relevant for each given analysis were concatenated row-wise to generate the versions used to fit the GLM models.

To evaluate the performance of the Ordinary Least Squares (OLS) models, trials were partitioned into training and test datasets, each containing 50% of the data. For the results shown in **Extended Data Fig. 3b** and **Extended Data Fig. 4c**, multiple model runs were carried out, with the number of repetitions designated *Y* in this paragraph. For each run, the data were split into training and test datasets and held constant for all the models tested in that run. *Y*=10 for the leave-out analysis (**Extended Data Fig. 3e**) and *Y*=3 for the hyperparameter analysis (**Extended Data Fig. 3b, Extended Data Fig. 4c**). For each model run, a 10-fold Group Shuffle Split (GSS) by trial was applied to the training set to obtain ranges for the mean squared errors (MSE), based on an 80-20 training-validation split within each of the 10 GSS folds. Each validation MSE value in the boxplots (**Extended Data Fig. 3c and 3d, Extended Data Fig. 4a and 4b**) is the average squared sum of the residuals across all validation datapoints in these 10 GSS folds. Finally, the model was refit to and evaluated on the entire training dataset, and this refit model was in turn evaluated on the test dataset, resulting in the MSEs and R^2^ values for each model run. The R^2^ values presented in the text are the average values calculated from the test sets averaged across *Y* model runs. Typically, these values had small variance, with ranges from maximum to minimum of <1.2% percentage points. Therefore, the ranges are not stated in the text.

For each of the Least Squares Regression models used, the algorithms minimize a cost function with respect to the fitted coefficients. The cost functions are as follows, where *J* is the cost function to be minimized, X is the design matrix, y is the response vector, *β* is the set of fitted coefficients, 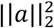 is the sum of the squared entries in vector a, ||*a*||_1_ is the sum of the absolute values of the entries in vector a, and *λ* is the L1 ratio.

*OLS*:

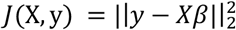

*Ridge Regression (L2):*

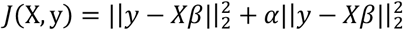

*Elastic Net / Lasso Regression (L1)*:

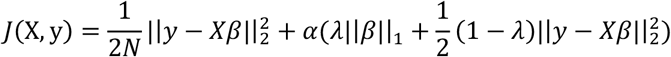

Note that for OLS, *α* = 0 as there is no regularization. Furthermore, setting *λ*=1 yields Lasso Regression (L1 Regularization). However, setting *λ*=0 does not give an equation equivalent to the version of Ridge Regression provided above. Instead, the residual term is divided by a factor 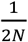 of relative to the Ridge Regression model described above resulting two different *α* scales (**Extended Fig. 3b** and **Extended Fig. 4c**). Additionally, for L2 Regularization, the validation-based models were fit to 80% of the total of samples available to the final model; thus, the validation models performed worse than their training or test counterparts because they are, in effect, facing an increased amount of regularization.

The sources for the Least Squares Regression models are listed below: OLS: https://scikitlearn.org/stable/modules/generated/sklearn.linear_model.LinearRegression.html L2: https://scikit-learn.org/stable/modules/generated/sklearn.linear_model.Ridge.html L1 & Elastic Net: https://scikit-learn.org/stable/modules/generated/sklearn.linear_model.ElasticNet.html

All kernels (*β* coefficients) depicted are the mean coefficients across the *Y* model runs with the standard error of the mean represented in the shaded regions. All GLM reconstructions depict the average signal with an overlay of the bootstrapped 95% confidence intervals of the upper and lower bounds (shaded region).

### Acute brain slice preparation

Brain slices were obtained from 2-to 4-month-old mice (both male and female) using standard techniques. Mice were anaesthetized by isoflurane inhalation and perfused cardiacly with ice-cold ACSF containing (in mM) 125 NaCl, 2.5 KCl, 25 NaHCO_3_, 2 CaCl_2_, 1 MgCl_2_, 1.25 NaH_2_PO_4_, and 25 glucose (295 mOsm kg^−1^). Brains were blocked and transferred into a slicing chamber containing ice-cold ACSF. Sagittal slices of striatum for amperometric or cell attached recordings were cut at 300 μm thickness with a Leica VT1000 s vibratome in ice-cold ACSF, transferred for 10 min to a holding chamber containing choline-based solution (consisting of (in mM): 110 choline chloride, 25 NaHCO_3_, 2.5 KCl, 7 MgCl_2_, 0.5 CaCl_2_, 1.25 NaH_2_PO_4_, 25 glucose, 11.6 ascorbic acid, and 3.1 pyruvic acid) at 34°C then transferred to a secondary holding chamber containing ACSF at 34C for 10 min and subsequently maintained at room temperature (20–22°C) until use. All recordings were obtained within 4 hours of slicing. Both choline solution and ACSF were constantly bubbled with 95% O_2_/5% CO_2_.

### Cell-attached recordings

Acute sagittal brain slices and electrophysiological recordings were obtained from the dorsal striatum as described before^75^, with the following variations: cholinergic interneurons were identified using morphological and electrophysiological features^22^. Slices were sustained in ACSF with 10 μM of gabazine, CPP, and NBQX (Tocris). For cell-attached recordings, bath temperature were maintained at 34C, pipettes were filled with ACSF, had 1-2MΩ resistance, seal resistances were from 10 to 100MΩ. Action potential firing was monitored in the cell-attached recording configuration in the voltage-clamp mode (V_hold_ = 0 mV). ChR2 was activated by a single 2 ms pulse of 473 nm light delivered at 5.74 mW using full-field illumination through the objective at 120 second intervals.

### Amperometry recordings

Slices were stimulated with 593 nm light, delivered at 5.86 mW for 2 ms using full-field illumination through the objective at 180 second intervals. Constant-potential amperometry was performed as previously described^75^. Briefly, glass-encased carbon-fiber microelectrodes (CFE1011 from Kation scientific - 7 μm diameter, 100 μm length) were placed approximately 50–100 μm within dorsal striatum slices and held at a constant voltage of + 600 mV for 9 second vs. Ag/AgCl by a Multiclamp 700B amplifier (Molecular Devices). Electrodes were calibrated with fresh 5 μM dopamine standards in ACSF to determine CFE sensitivity and to allow conversion of current amplitude to extracellular dopamine concentration.

## Acknowledgements

We thank all members of the Sabatini lab for helpful experimental suggestions and advice, in particular M.L. Wallace and S.J. Lee. We thank P. Capelli, A.E. Girasole, and C.S. Smillie for feedback on the manuscript. We also thank G. Radeljic for technical assistance and Yulong Li for generously sharing the rDAh sensor. We are deeply grateful to Ofer Mazor and Pavel Gorelick at the Harvard Medical School Research Instrumentation Core Facility for their help with implementing the hardware and software for the 2ABT. This work was supported by grants to B.L.S (R37-NS046579, U19 NS113201, and Simons Center for the Global Brain), to L.C. (Hanna Gray Fellowship from the Howard Hughes Medical Institute), and to C.C.B. (NSF Graduate Research Fellowship Program).

## Author contributions

L.C. and B.L.S. conceptualized the study. L.C. and B.L.S. wrote and edited the manuscript, with feedback from C.C.B. and J.A.Z. L.C. and M.J.W. performed experiments except electrophysiological ones, which were done by W.W. B.L.S. developed the photometry analysis pipeline, and C.C.B. established the behavioral analysis pipeline. J.A.Z. and B.L.S conceptualized and implemented the GLMs.

## Data and code availability

The data and code that supports the findings of this study are available upon request from the corresponding author.

---

**Extended Data Fig. 1.**
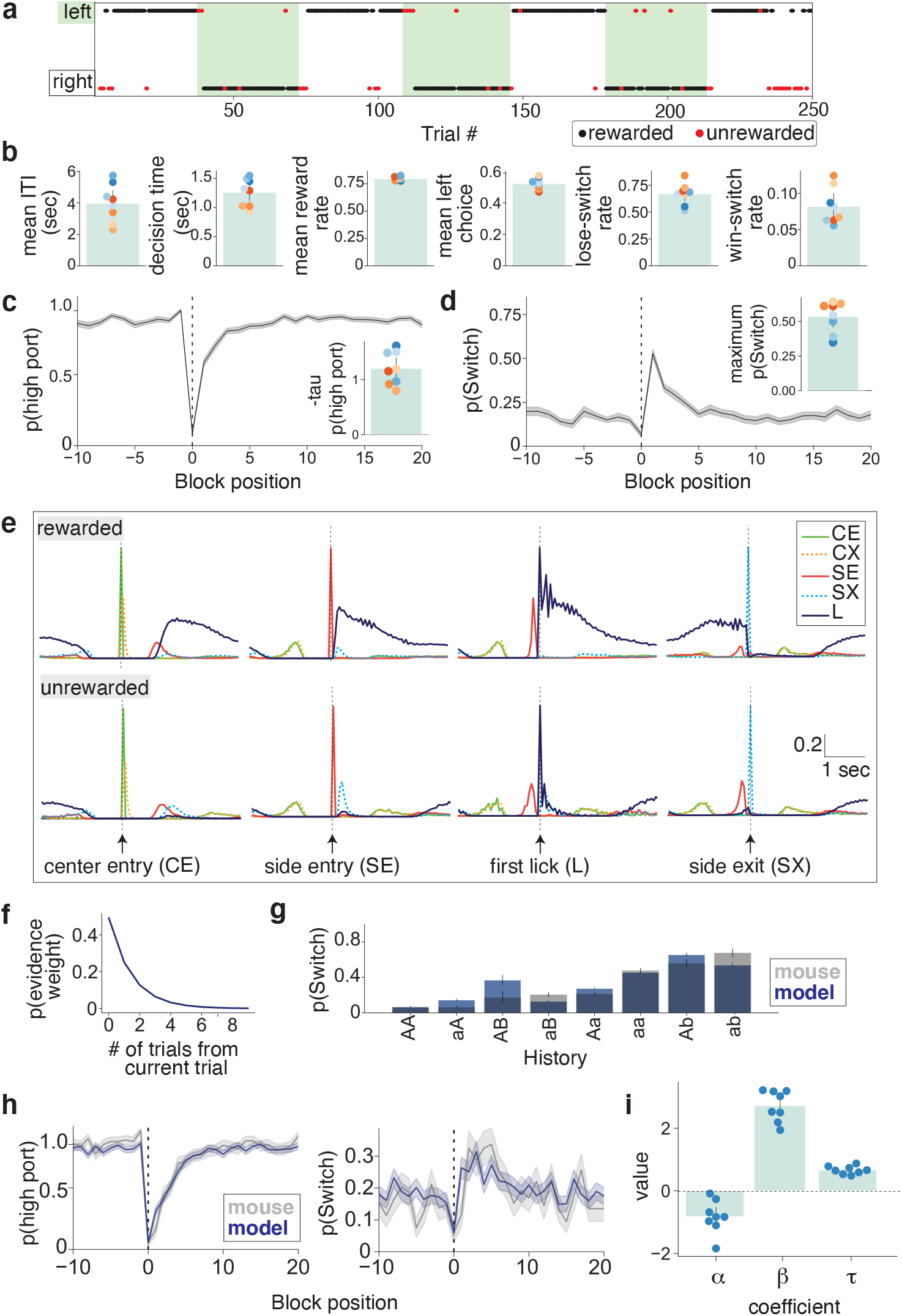
Baseline behavior and behavioral modeling of mouse performance in the 2ABT. **a**. Baseline expert behavior during a 2ABT session for a representative mouse. Green and white shading denote the left port and right port, respectively, as being the higher rewarded port. The location of dots represents a choice made by the mouse. A choice resulting in a rewarded outcome is indicated with a black dot while a red dot represents an unrewarded trial. **b**. The average intertrial interval (ITI) and decision time, and the rate of rewarded trials, left choice, and switching following an unrewarded (lose-switch) or rewarded (win-switch) outcome on the previous trial for a representative group of mice. Each colored dot represents a unique mouse (n = 7). Error bars denote the 95% confidence interval. **c**. Probability of occupancy at the highly rewarded port (p(high port)) as a function of block position. The average is indicated with a bolded line and the standard error of the mean (SEM) is overlaid in the shaded region. Calculated taus are shown (inset), with each dot representing a unique mouse and error bars denoting the 95% confidence interval. Tau is the time constant of recovery to the higher rewarded port, in units of trial number. **d**. Probability of switching ports (p(Switch)) as a function of block position and the maximum probability of switching (inset), which is defined across all block positions. Data is shown as in (**c**). **e**. Average timing of behavioral events in rewarded and unrewarded trials. The data are aligned to the indicated behavioral events: center port entry (CE), center port exit (CX), side port entry (SE), side port exit (SX), and first lick (L) (n = 7 mice). **f**. Evidence weights as a function of the indicated number of trials from the current one. Evidence weights are defined as the proportion of evidence that each trial contributes to the mouse’s action and reward history, calculated based on the average *τ* coefficient in (**i**). **g**. Conditional switch probabilities for each action-outcome trial sequence of history length two, sorted by increasing switch probability. The original data (gray) is overlaid with the data predicted by the recursively formulated logistic regression RFLR model (blue). The bar heights show the mean switch probability across mice for each corresponding sequence history, and the error bars depict the binomial standard error for the mouse test data. **h**. RFLR predicted probability (blue) of choosing the highly rewarded probability port (left) and of switching ports (right) as a function of trial number from the block transition at zero, as compared to the mouse behavior (gray). The mean across trials (bold line) at the same block position and the standard error (shaded region) are shown. **i**. Summary of the *α, β* and *τ* coefficients from the RFLR model. Each blue dot represents an individual mouse. Error bars denote the 95% confidence interval.

**Extended Data Fig. 2.**
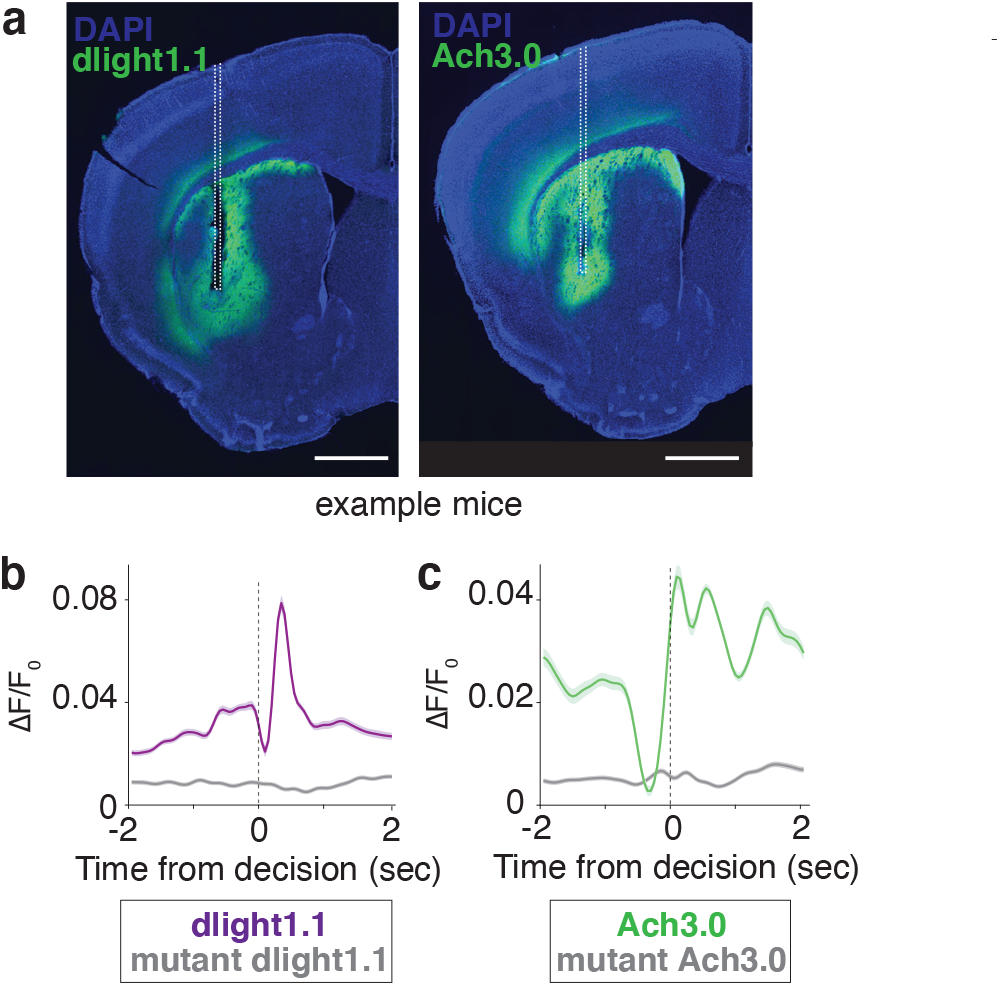
Histology and photometry controls for DA and Ach recordings in the VLS. **a**. Histology depicting dlight1.1 expression (left) and Ach3.0 expression (right) in representative mice. The optical fiber tract is indicated with the dashed white line. Scale bar: 1 mm. **b**. Average ΔF/F_0_ of DA release during rewarded trials from dlight1.1 or the DA-binding mutant of dlight1.1 (n = 4 mice). Signals (bold line) are aligned to side port entry with the standard error overlaid (shaded region). **c**. Average ΔF/F_0_ of Ach release during rewarded trials from Ach3.0 or the mutant version of Ach3.0 (n = 4 mice). Signals are depicted as in (**b**).

**Extended Data Fig. 3.**
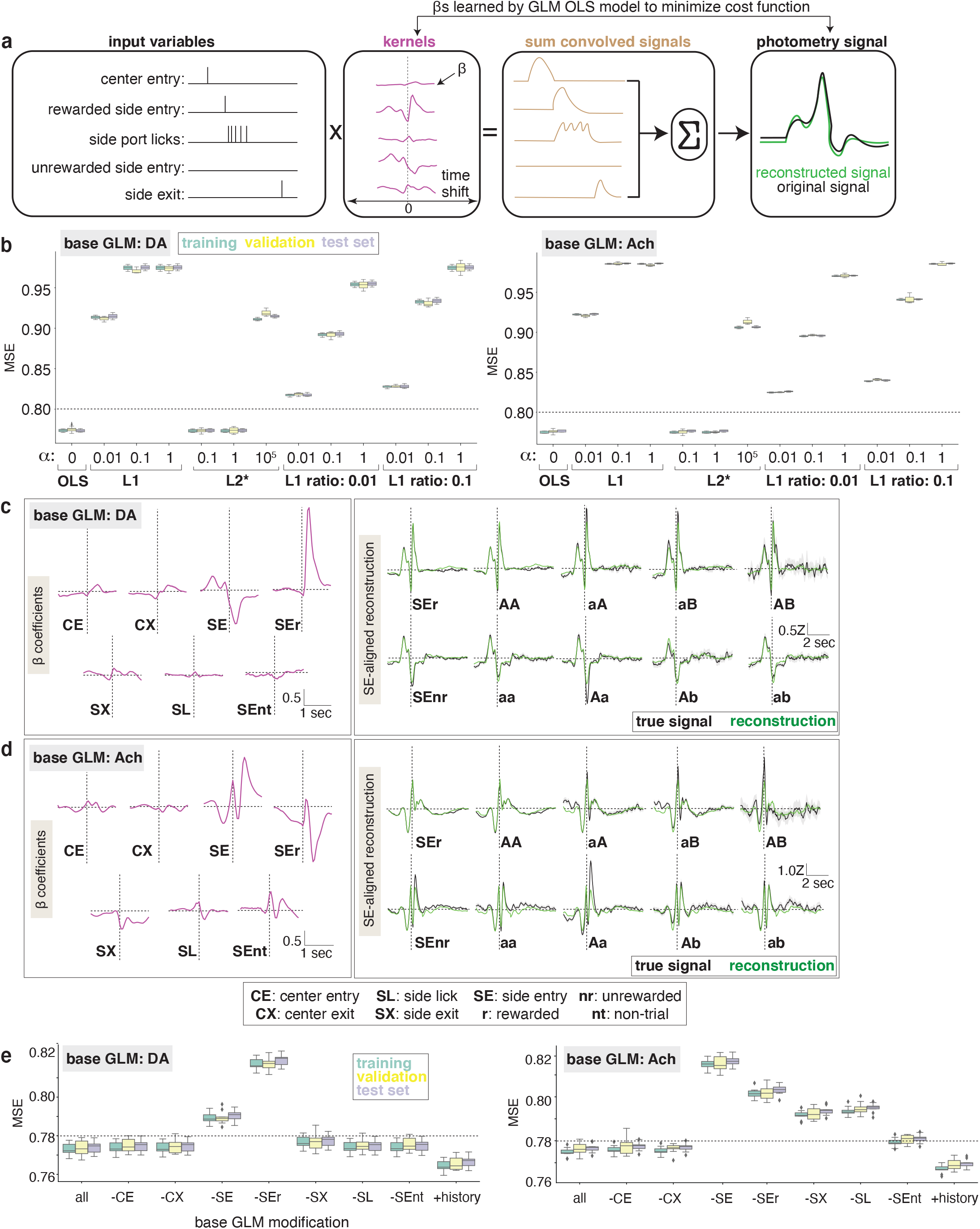
General linear models (GLMs) of DA and Ach with a base behavior feature set. **a**. Schematic of the GLM workflow. A set of input variables are convolved with their associated time-based kernels, with each time step consisting of a separate *β* coefficient fit by minimizing an ordinary least squares (OLS) cost function. The resulting convolved signals are summed to generate the reconstructed photometry signal. **b**. The effect of different hyperparameter sweeps over models associated with Least Squares Regression – OLS, Lasso Regression (L1), Ridge Regression (L2), and Elastic Net (L1+L2) on the training, validation, and test set mean squared errors (MSE) of the DA (left) and Ach (right) base GLMs. **c**. DA kernels (left) and side entry (SE) aligned reconstructed and true DA signals (right) for the base GLM model. The average signal (bold line) and 95% confidence interval (shaded region) are depicted (n = 8 mice). **d**. Ach kernels (left) and SE-aligned reconstructed and true Ach signals (right) for the base GLM model. Data is depicted as in (**b**) (n = 9 mice). **e**. The effect of omission (-) or inclusion (+) of the indicated input variables on the DA (left) and Ach (right) base GLM performance, as measured by the effect on training, validation, and test set MSE.

**Extended Data Fig. 4.**
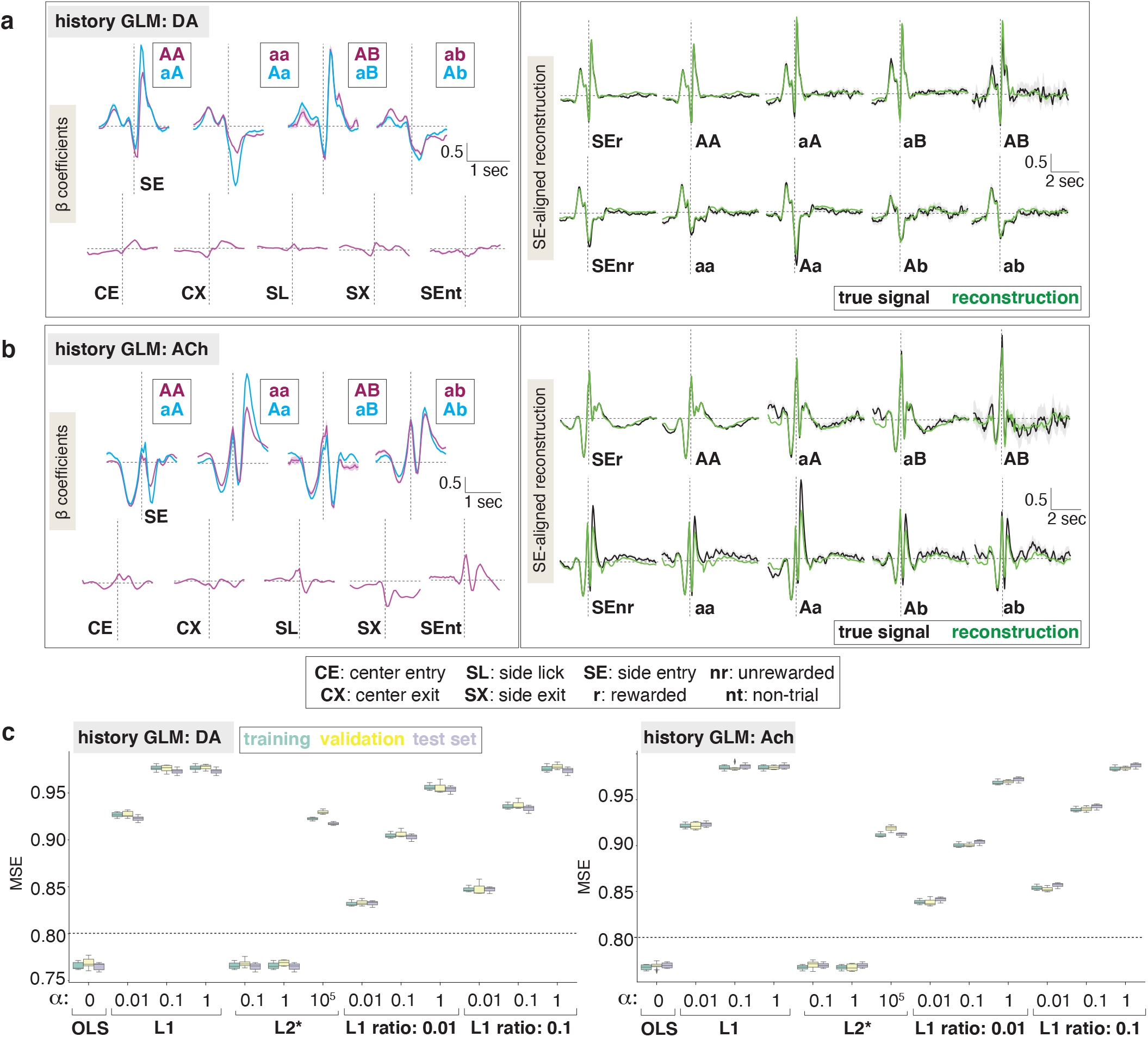
DA and Ach GLMs incorporating a history feature set. **a**. DA kernels (left) and side entry (SE) aligned reconstructed and true DA signals (right) for the history GLM model. The average signal (bold line) and 95% confidence interval (shaded region) are depicted (n = 8 mice). **b**. Ach kernels (left) and SE-aligned reconstructed and true Ach signals (right) for the history GLM model. Data is depicted as in (**a**) (n = 9 mice). **c**. The effect of different hyperparameter sweeps over models associated with Least Squares Regression – OLS, Lasso Regression (L1), Ridge Regression (L2), and Elastic Net (L1+L2) on the training, validation, and test set mean squared errors (MSE) of the DA (left) and Ach (right) history GLMs.

**Extended Data Fig. 5.**
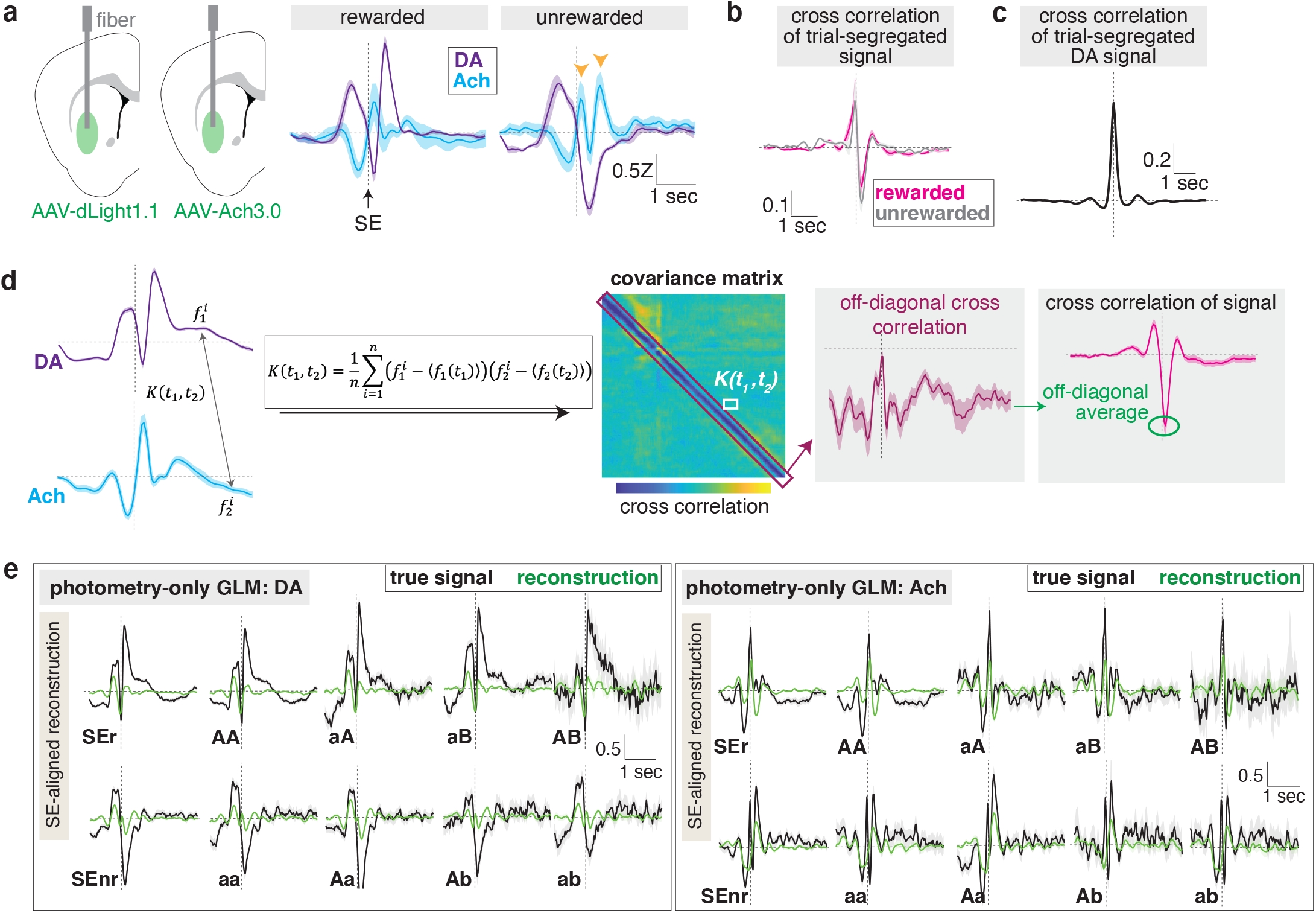
Cross correlation and GLM analyses of DA and Ach signals. **a**. Injection and implantation scheme for VLS recordings of DA and Ach release from separate mice (left). The average Z score of the sensor signal (bold line) is aligned to side port entry (SE) (middle and right) with the standard error (SEM) overlaid (shaded region) (DA: n = 7; Ach: n = 9 mice). Orange arrows denote the double rise of Ach referenced within the main text. **b**. Cross correlation of trial segregated DA and Ach signals recorded from mice in (A) for rewarded and unrewarded trials, in which DA lags Ach. The average cross correlation is shown (bold line) with the standard error indicated (shaded region). **c**. Cross correlation of trial segregated DA signals recorded with dlight1.1 from opposite hemispheres of the same brain (n = 4 mice). **d**. Summary of the correlation analyses in **Fig. 2**. A covariance matrix is built by calculating the cross correlation (*K*(*t*_1_, *t*_2_)) of the DA signal at one time point 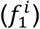 to the Ach signal at all other time points (one time point is shown as 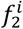), and vice versa, using the indicated equation where ⟨*f* (*t*_1_)⟩ is the mean across all trials for a certain time point, *t*. In this covariance matrix, the off-diagonal signal shows a striking negative cross correlation. The average of this signal is equivalent to the minimum cross-correlation value calculated from the trial-averaged signals. **e**. Side entry (SE)-aligned reconstructed and true DA and Ach signals from GLMs that only incorporate a photometry variable. The average signal (bold line) and the 95% confidence intervals (shaded region) are shown.

**Extended Data Fig. 6.**
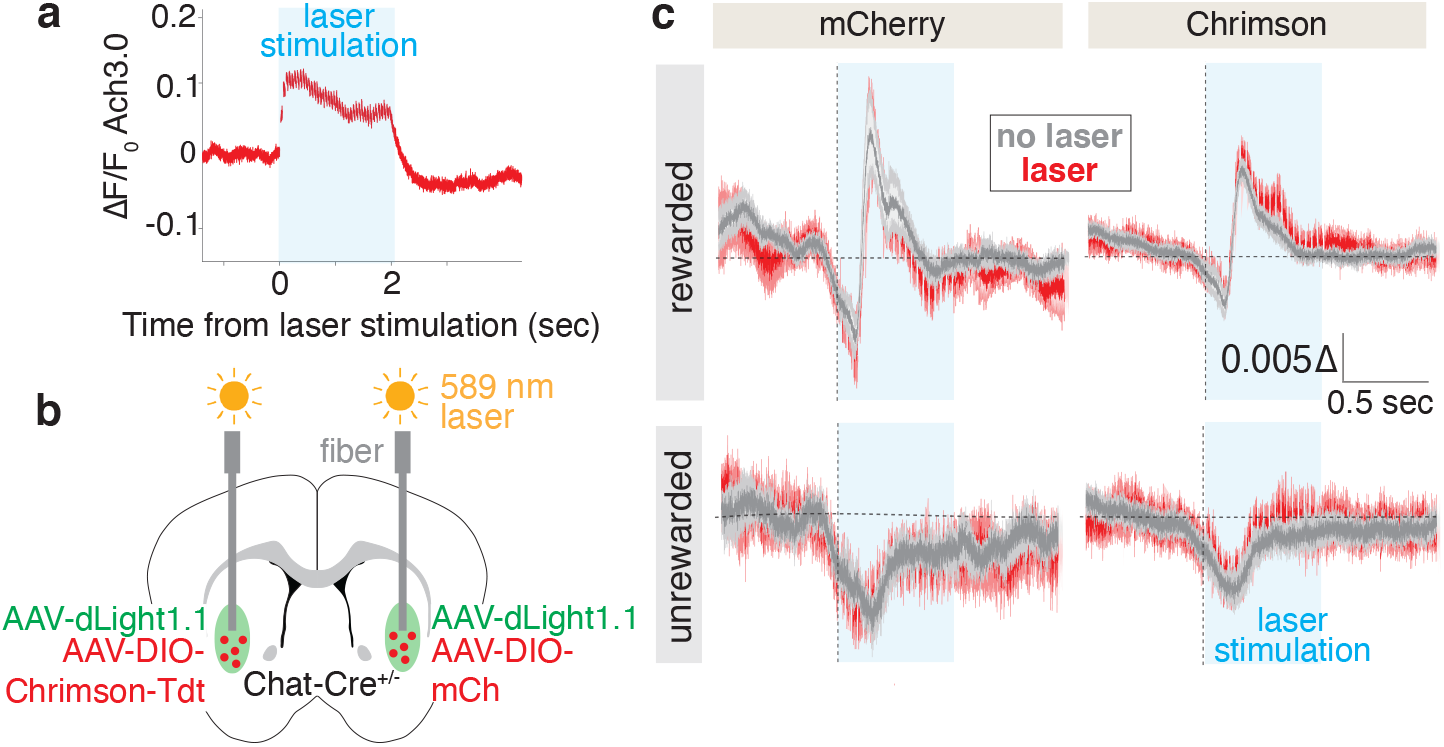
Optogenetic perturbations of CINs in the VLS. **a**. Ach release upon optogenetic activation of Chrimson-expressing CINs. Average ΔF/F_0_ of Ach3.0 is depicted (bold line) with standard error (SEM) overlaid (red shaded region) and the laser stimulation artifacts omitted. The duration of the laser stimulation is shown (blue shaded region). **b**. Injection and optical fiber implantation for animals recorded in (**c**). dLight1.1 is coexpressed in VLS with either Chrimson or mCherry (mCh) expressed in CINs of separate hemispheres. **c**. Optogenetic activation during the indicated trial types of CINs expressing Chrimson or mCherry. Average ΔF/F_0_ (bold line) is depicted for trials with and without laser stimulation, aligned to side port entry (dashed vertical line), with SEM overlaid (red and gray shaded regions) and the laser stimulation artifacts omitted (n = 7 mice).

**Extended Data Fig. 7.**
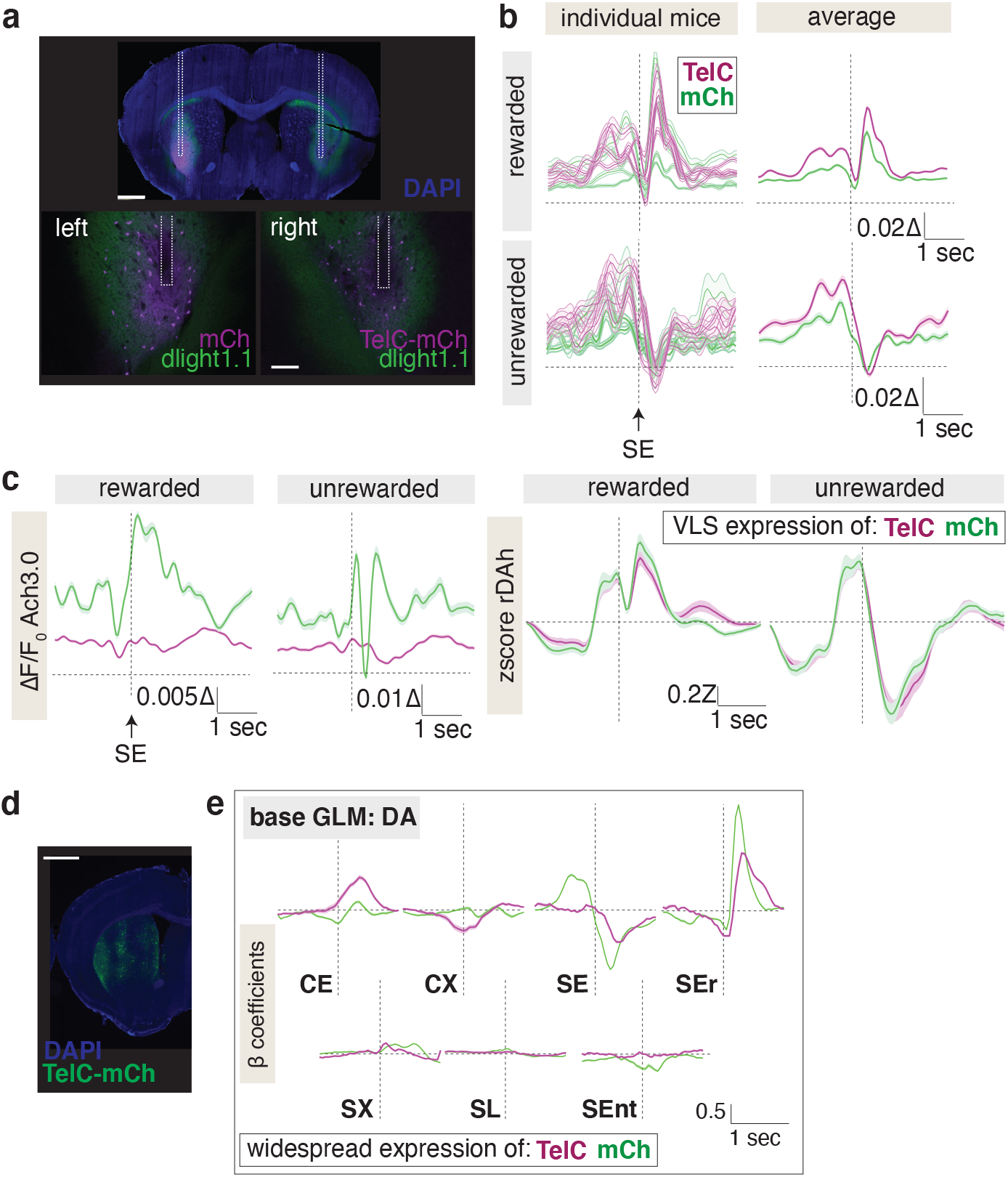
Histology, photometry and GLM analyses of the effects of CIN-specific tetanus expression on striatal DA signals. **a**. Histology of a representative mouse with dlight1.1 expression in both hemispheres and mCherry (mCh) or tetanus (TelC) expression in CINs of separate hemispheres. Fiber tract is outlined with white dashed lines. Scale bar: 1 mm. **b**. Effect of TelC or mCh expression in CINs on DA release. Side-entry (SE) aligned ΔF/F_0_ signals for individual mice (left) and the average ΔF/F_0_ (bold line) across all mice of a treatment group (right) are depicted with standard error overlaid (shaded region) (n = 4 mice). **c**. Simultaneous recordings of Ach (left) and DA (right) signals from mice with CINs in the VLS expressing TelC or mCh. Data is represented as in (**c**) (n = 4 mice). **d**. Histology from a representative mouse with TelC expression in CINs of the entire striatum. Scale bar: 1 mm. **e**. DA kernels for the base GLM derived from recordings in **Fig. 3f** with striatum-wide expression of the indicated proteins in CINs. The average signal (bold line) and the confidence interval (shaded region) are shown.

**Extended Data Fig. 8.**
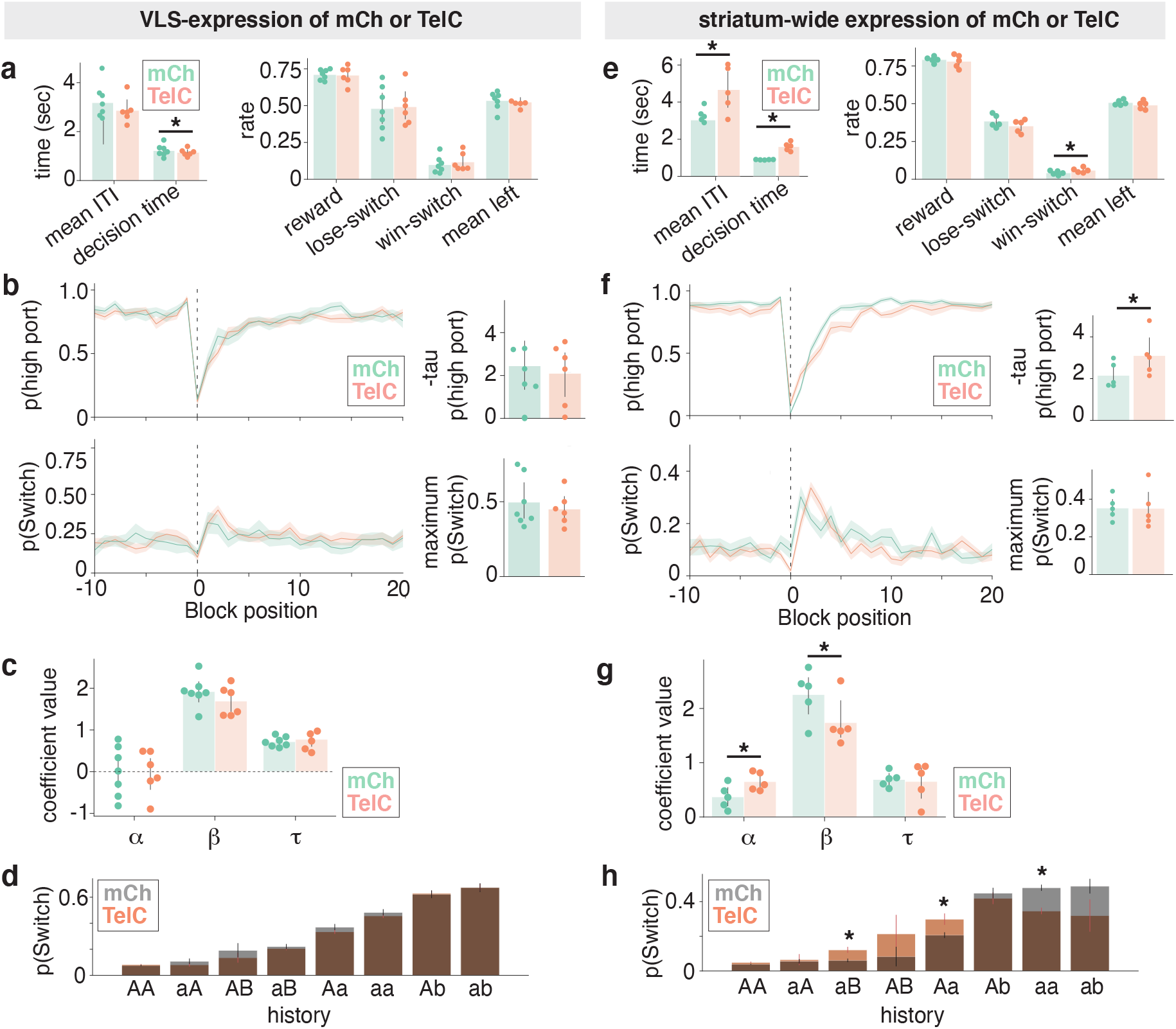
Behavioral analysis of mice with VLS-specific or striatal-wide tetanus expression in CINs. **a**. Indicated performance metrics for mice from **Fig. 3e** that express tetanus toxin (TelC) or mCherry (mCh) control protein in CINs of the VLS. Each dot represents a unique mouse. Error bars denote the 95% confidence interval, and significance is determined as >95% confidence from bootstrapped samples (asterisk). **b**. Probability of occupancy at highly rewarded port (p(high port)) or switching (p(Switch)) as a function of block position for mice from **Fig. 3e** (left). The average is indicated with a bold line and the standard error of the mean (SEM) is overlaid in the shaded region. Calculated taus and maximum pSwitch rates (right) are shown, with each dot representing a unique mouse. Error bars denote the 95% confidence interval. **c**. Summary of the *α, β* and *τ* coefficients generated from the RFLR model of the indicated treatment groups for mice from **Fig. 3e**. Data is depicted as in (**a**). **d**. Conditional switch probabilities for the indicated action-outcome trial sequence sorted by increasing switch probability for mice from **Fig. 3e**. Mice with CINs expressing mCh (gray) are overlaid with those expressing TelC (orange). The bar heights show the mean switch probability across mice for each corresponding sequence history, and the error bars depict the binomial standard error for the mouse test data. **e**. The indicated performance metrics for mice from **Fig. 3f** with striatum-wide CIN expression of TelC or mCh. Data is depicted as in (**a**). **f**. Probability of occupancy at highly rewarded port (p(high port)) or switching (p(Switch)) as a function of block position for mice from **Fig. 3f** (left). Data is depicted as in (**b**). **g**. Summary of the *α, β* and *τ* coefficients generated from the RFLR model for mice from **Fig. 3f**. Data is depicted as in (**c**). **h**. Conditional switch probabilities for mice from **Fig. 3f**. Data is depicted as in (**d**).

**Extended Data Fig. 9.**
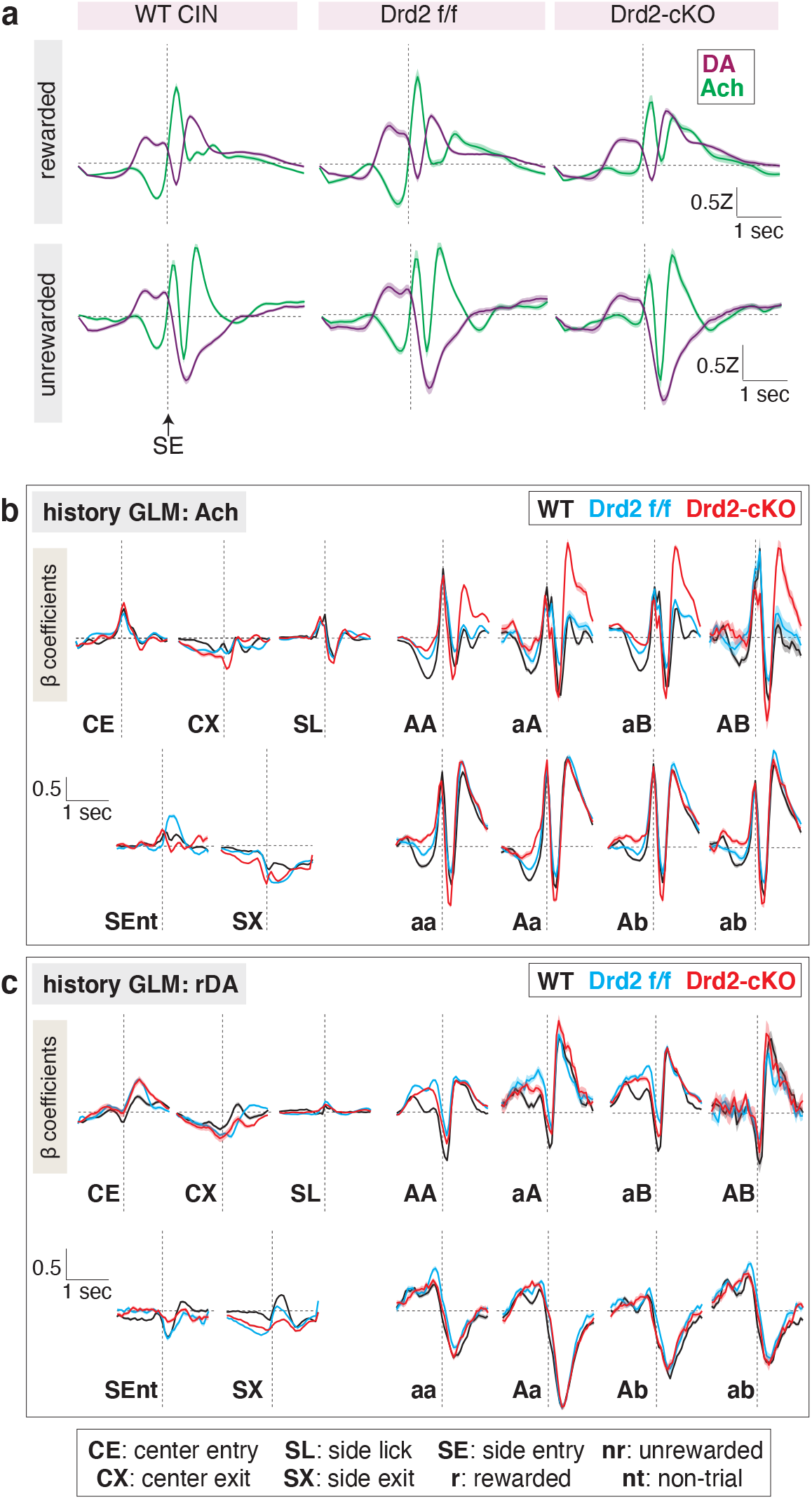
Photometry and GLM analyses of DA and Ach signals from mice lacking D2R expression in CINs. **a**. Overlay of Ach and DA release aligned to side port entry (SE) recorded from mice in **Fig. 4g**. Average release is depicted (bold line) with standard error of the mean (SEM) overlaid (shaded region) (WT: n = 12 mice; Drd2 f/f: n = 7 mice; Drd2-cKO: n = 8 mice). **b**. Kernels of the indicated input variables for the history-based GLM of Ach signals from mice in **Fig. 4g**. Average *β* coefficients are depicted (bold line) with confidence interval outlined (shaded region). **c**. Kernels of the indicated input variables for the history-based GLM of rDAh signals from mice in **Fig. 4g**. Data is displayed as in (**b**).

**Extended Data Fig. 10.**
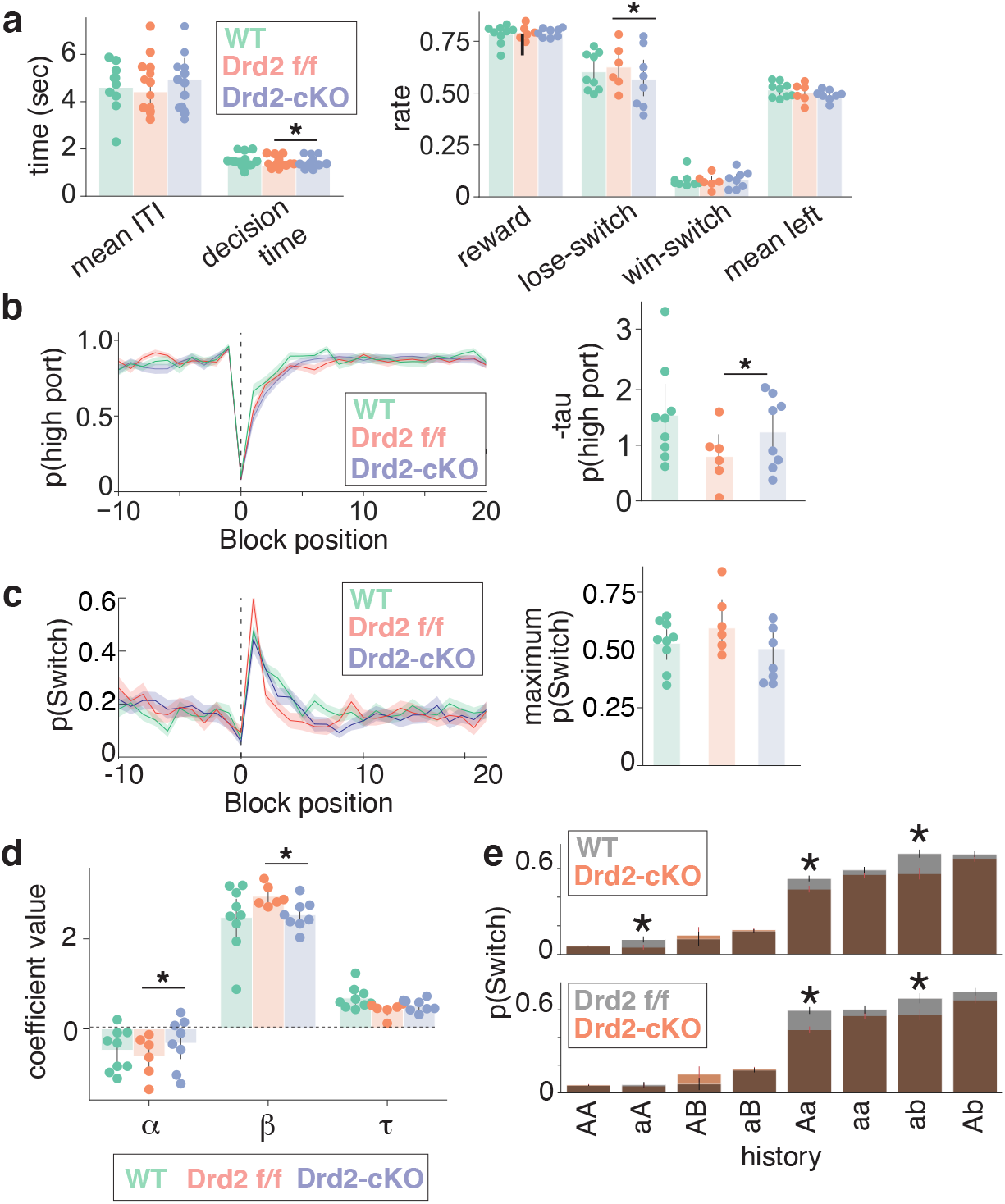
Behavioral effects of mice lacking D2R expression in CINs. **a**. Indicated performance metrics for mice in **Fig. 4g**. Each dot represents a unique mouse with binomial error overlaid. Error bars denote the 95% confidence interval, and significance is denoted as >95% confidence from bootstrapped samples (asterisk). **b**. Probability of occupancy at highly rewarded port (p(high port)) as a function of block position for mice in **Fig. 4g** (left). The average is indicated with a bold line and the SEM is overlaid in the shaded region. Calculated taus (right) are shown, with each dot representing a unique mouse and error bars depicting the 95% confidence interval. **c**. Probability of switching (p(Switch)) as a function of block position for mice in **Fig. 4g** (left). Data is depicted as in (**b**). Calculated maximum switch rates (right) are shown, with each dot representing a unique mouse and error bars depicting the 95% confidence interval. **d**. Summary of the *α, β* and *τ* coefficients generated from the RFLR model of the indicated mice in **Fig. 4e**. Each dot represents an individual mouse and error bars show the 95% confidence interval. **e**. Conditional switch probabilities for the indicated action-outcome trial sequence sorted by increasing switch probability for mice from **Fig. 4e**. The bar heights show the mean switch probability across mice for each corresponding sequence history, and the error bars depict the binomial standard error. Significance is denoted as >95% confidence from bootstrapped samples (asterisk).

**Extended Data Fig. 11.**
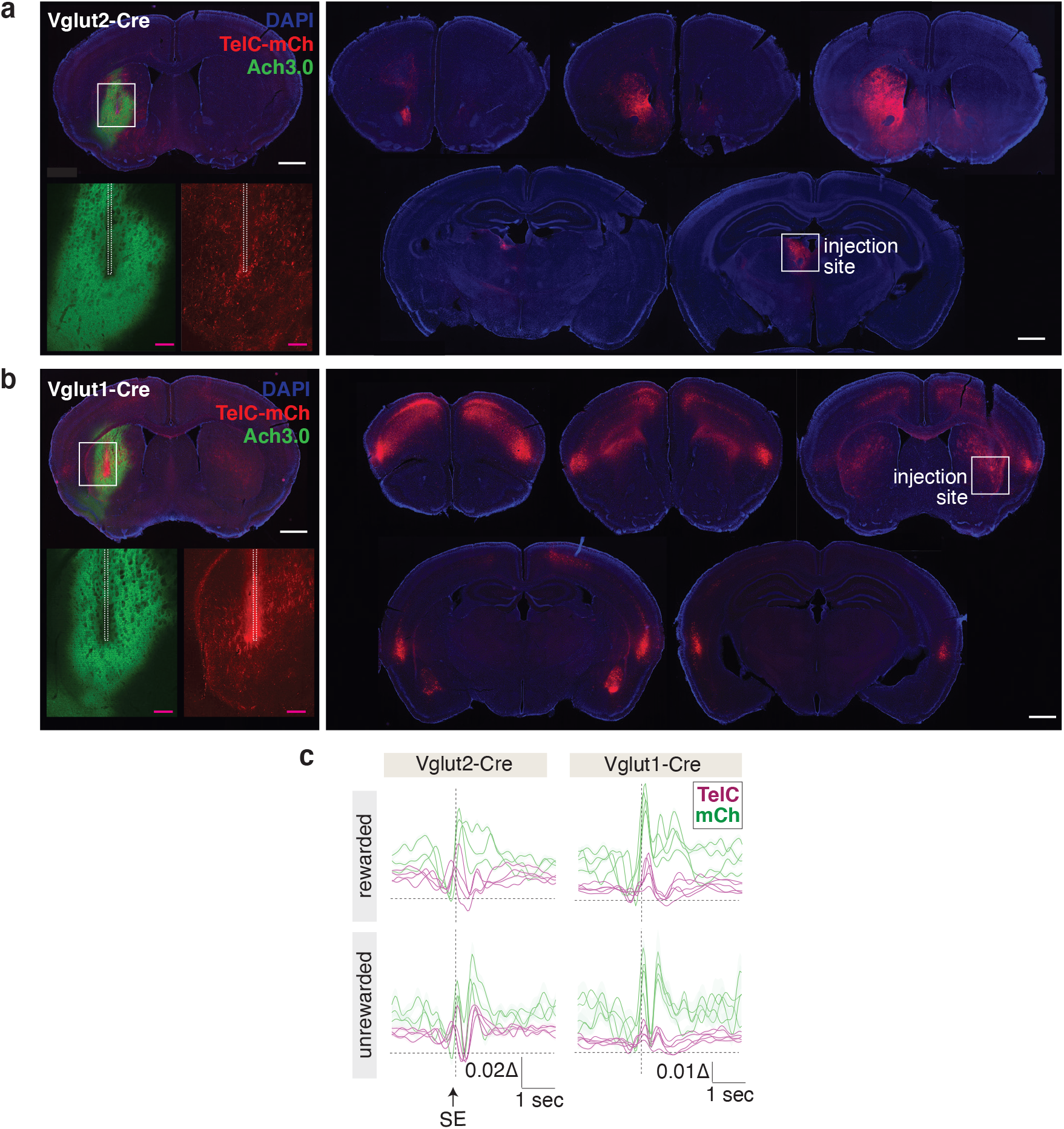
Histology and photometry analyses of tetanus toxin expression in cortical and thalamic inputs. **a**. Histology of a representative mouse expressing tetanus toxin (TelC-mCh) in thalamus. Epifluorescence image of a coronal slice containing the fiber implantation site (top left) with a magnified view of the Ach3.0 and TelC expression sites (bottom left insets). Coronal slices spanning the anterior to posterior regions of the brain showing the injection site in thalamus (right) and TelC-positive terminals projecting to striatum. Scale bar (white): 1 mm; Scale bar (pink): 200 μm. **b**. Example histology of a representative mouse expressing TelC toxin in cortex. Images are depicted as in (**a**). **c**. Effect of Cre-dependent TelC or mCh expression in Vglut2-Cre (left) or Vglut1-Cre (right) mice on Ach release, as measured by average ΔF/F_0_ for individual mice (bold line) aligned to side port entry (SE) with standard error (shaded region) overlaid.

**Extended Data Fig. 12.**
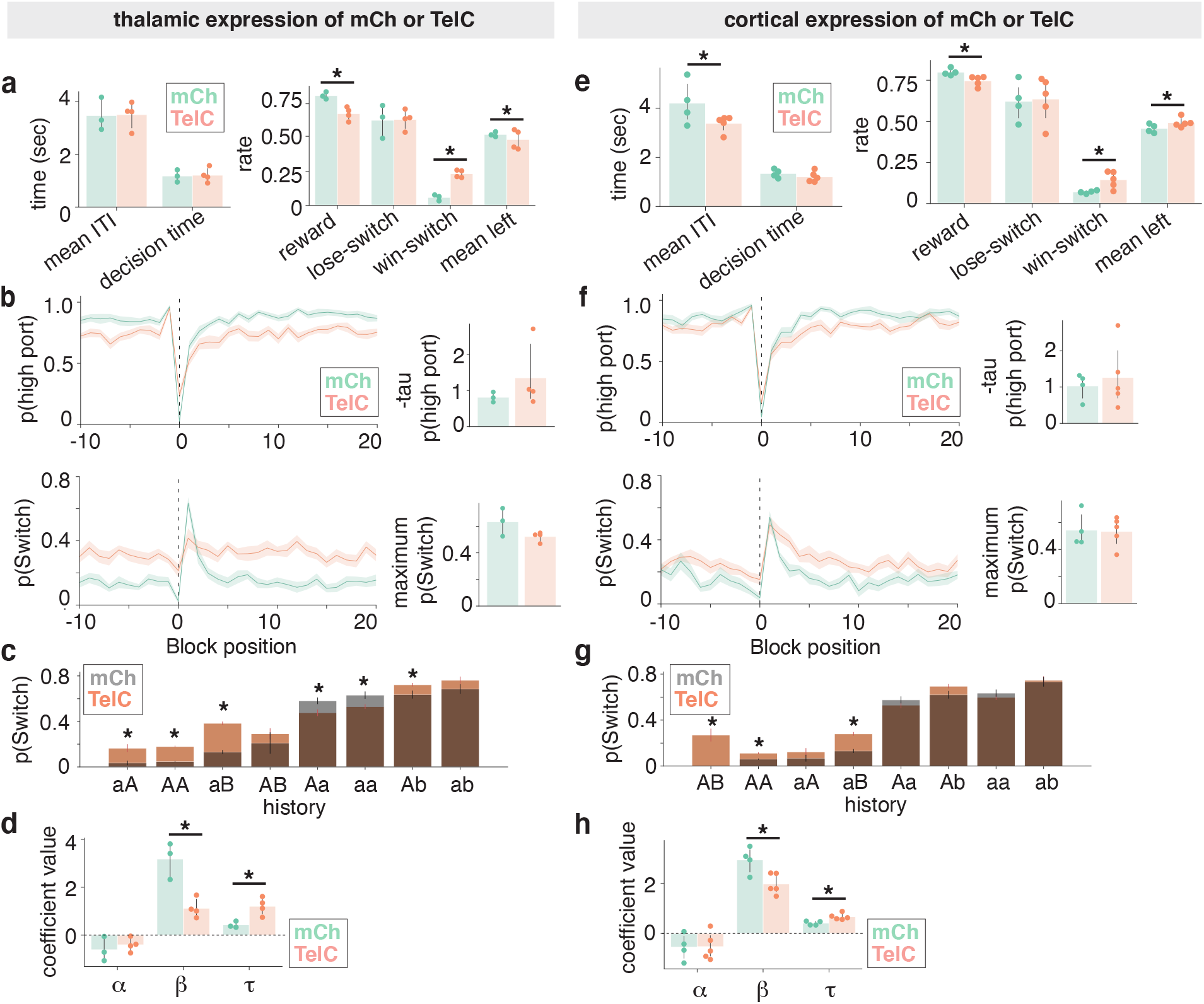
Behavioral effects of perturbing cortical and thalamic inputs into the VLS. **a**. Indicated performance metrics for mice in **Fig. 5e** that express tetanus toxin (TelC) or mCherry (mCh) control protein in the thalamus. Each dot represents a unique mouse. Error bars depict the 95% confidence interval, and significance is determined as >95% confidence from bootstrapped samples (asterisk). **b**. Probability of occupancy at highly rewarded port (p(high port)) or switching (p(Switch)) as a function of block position (left) for mice in **Fig. 5e**. The average is indicated with a bold line and the SEM is overlaid in the shaded region. Calculated taus and maximum pSwitch rates (right) are shown, with each dot representing a unique mouse. Error bars depict the 95% confidence interval. **c**. Conditional switch probabilities for the indicated action-outcome trial sequences sorted by increasing switch probability for mice in **Fig. 5e**. Mice with CINs expressing mCh (gray) are overlaid with those expressing TelC (orange). The bar heights show the mean switch probability across mice for each corresponding sequence history, and the error bars depict the binomial standard error. Significance is determined as >95% confidence from bootstrapped samples (asterisk). **d**. Summary of the *α, β* and *τ* coefficients generated from the RFLR model for mice in **Fig. 5e**. Data is depicted as in (**a**). **e**. The indicated performance metrics for mice in **Fig. 5f** that express TelC or mCh in the VLS-specific cortical inputs. Data is depicted as in (**a**). **f**. Probability of occupancy at highly rewarded port (p(high port)) or switching (p(Switch)) as a function of block position for mice in **Fig. 5f**. Data is depicted as in (**b**). **g**. Conditional switch probabilities of the indicated treatment groups for mice in **Fig. 5f**. Data is depicted as in (**c**). **h**. Summary of the *α, β* and *τ* coefficients generated from the RFLR model of the indicated treatment groups in **Fig. 5f**. Data is depicted as in (**d**).

